# Characterization of Microbubble Cavitation in Theranostic Ultrasound-mediated Blood-Brain Barrier Opening and Gene Delivery

**DOI:** 10.1101/2025.01.17.633644

**Authors:** Fotios N. Tsitsos, Alec J. Batts, Daniella A. Jimenez, Chunqi Li, Robin Ji, Sua Bae, Angeliki Theodorou, Samantha L. Gorman, Elisa E. Konofagou

**Affiliations:** Department of Biomedical Engineering, Columbia University, New York, NY, USA; Department of Radiology, Columbia University, New York, NY, USA; Department of Neurological Surgery, Columbia University, New York, NY, USA

**Keywords:** Theranostic ultrasound, short pulses, microbubbles, cavitation, viral delivery

## Abstract

**Rationale:** The characterization of microbubble activity has proven critical in assessing the safety and efficacy of ultrasound-mediated blood-brain barrier (BBB) opening and drug and gene delivery. In this study, we build upon our previous work on theranostic ultrasound (ThUS)-mediated BBB opening (ThUS-BBBO) and conduct for the first time a comprehensive characterization of the role of microbubble cavitation in ThUS-BBBO, as well as its impact on gene delivery with adeno-associated viruses (AAV).

**Methods:** A repurposed imaging phased array was used throughout the study to generate focused transmits and record microbubble activity through high-resolution power cavitation imaging (PCI). The cavitation of microbubbles under ThUS pulses was first characterized in flow phantom using pulse lengths ranging from 1.5 to 20 cycles and under varying microbubble flow rates using a separate single-element transducer a passive cavitation detector (PCD). A comprehensive *in vivo* study in mice was then conducted to characterize the *in vivo* microbubble activity under ThUS and correlate the resulting cavitation with AAV-mediated transgene delivery and expression. The transcranial microbubble activity was first detected in two mice using a PCD, to confirm the findings of the flow phantom study. Next, three mouse studies were conducted to evaluate the relationship between cavitation and AAV delivery; one with three different microbubble size distributions using polydisperse and size-isolated microbubbles, one with variable burst length and burst repetition frequency, and one with different AAV serotypes and injection doses. Electronic beam steering was used to induce bilateral BBB opening with 1.5 cycle on the left and 10 cycles on the right hemisphere. Cavitation dose was correlated with BBB opening volume, AAV transgene expression was evaluated with immunofluorescence staining and histological safety was assessed with T2* imaging and Hematoxylin and Eosin staining.

**Results:** Frequency domain analysis in the phantoms revealed a broadband-cavitation dominance at the shorter pulse lengths, while harmonic cavitation components are significantly increased for longer pulses. The PCD was better at detecting higher frequency harmonics, while the signal received by the theranostic array was more broadband dominated. Analysis of signals in the time domain showed that the longer pulses induce higher microbubble collapse compared to short pulses. In the transcranial *in vivo* experiments, the PCD was able to detect increased harmonic cavitation for 10-cycle pulses. The microbubble study showed that 3-5 μm microbubbles resulted in the largest cavitation doses, BBBO volumes and AAV transgene expression compared to the smaller microbubble sizes. The burst sequence study revealed that the sequences with shorter bursts and faster burst repetition frequencies induce larger BBBO volumes and AAV transduction due to faster microbubble replenishment in the focal volume. Increased erythrocyte extravasation was observed on the hemisphere sonicated with 10-cycle pulses. Transgene expression was also increased with injection dose, without notable side effects during the three-week survival period. Finally, AAV9 was shown to be the serotype with the highest transduction efficiency compared to AAV2 and AAV5 at the same injected dose.

**Conclusions:** This is the first comprehensive study into the microbubble cavitation under theranostic ultrasound. The phantom and *in vivo* studies show that the mechanism of ThUS-BBBO is mainly transient cavitation dominant, as microbubble collapse increases with pulse length despite the increased harmonic frequency response. Increased cavitation dose resulted in larger BBBO volumes and transgene expression *in vivo*. While ThUS induced microhemorrhage for most of the studied conditions, it did not have an impact on the survival and behavior of the mice.

## Introduction

The effective treatment of central nervous system (CNS) diseases is often hindered by the presence of the blood-brain barrier (BBB), a specialized system of endothelial cells and tight-junction proteins lining the walls of capillary blood vessels in the brain and blocking the exchange of molecules larger than 400 Da between circulation and the brain parenchyma [1], [2]. Currently, efforts to increase the bioavailability of therapeutics in the neuronal space rely on pharmacological adaptations, which require complex chemical processes and show mild success [3], [4], or invasive surgical injections [5], which are often unfavorable for patients and suffer from potential infection complications and backflow limitations. To date, focused ultrasound (FUS), in combination with systemically administered microbubbles (MB), has been the only safe, non-invasive, and transient method for opening the BBB, allowing enhanced exchange of otherwise impermeable molecules through the barrier [6], [7], [8]. FUS-mediated BBB opening (FUS-BBBO) has been primarily studied for drug and gene delivery [9], [10], [11], [12]; however, it has also shown promise as a method of reversibly modulating the neuro-immune system [13], [14], [15] and enhancing disease biomarker detection through liquid biopsy [16], [17], [18], [19], [20], [21].

The mechanism by which FUS safely and locally opens the BBB is MB cavitation, which refers to the periodic oscillation of gas-filled MBs within the focal volume of the FUS beam. Cavitating MBs exert mechanical forces upon the capillary walls, enhancing transcellular and paracellular diffusion of large molecules through the BBB [22], [23]. A wealth of literature exists on the characterization of MB activity under various conditions of FUS. Generally, MB cavitation is classified as stable or inertial; stable cavitation refers to the sustained expansion and micro-streaming of MBs over time and is observed as enhanced secondary acoustic emissions at harmonics, sub-, and ultraharmonics of the fundamental frequency of sonication, whereas inertial or transient cavitation is characterized by microbubble collapse and production of high-velocity jets, manifesting as broadband acoustic signal in the frequency domain [24], [25], [26]. Studies have shown that safe and effective FUS-BBBO is achieved with parameters inducing primarily stable cavitation, as excessive inertial cavitation is typically associated with increased likelihood of tissue damage [24], [27]. Therefore, accurate cavitation monitoring is required to ensure that cavitation doses stay within the stable regime. An additional criterion to assess the risk of adverse cavitation-related bioeffects is the mechanical index (MI), which is defined as the ratio between the derated peak rarefactional pressure and the square-root of the fundamental frequency of sonication [28]. In the context of BBB opening, a mechanical index over 0.5 is typically associated with the onset of inertial-cavitation-related tissue damage, and thus most pre-clinical and clinical setups choose a combination of pressure and frequency that induces an MI below this threshold [29], [30].

Cavitation monitoring is conventionally achieved using a separate transducer, which passively receives acoustic emissions as microbubbles are being engaged by a geometrically focused transducer. Theranostic ultrasound (ThUS) was developed as a simplified and versatile tool for BBBO to achieve combined treatment and cavitation monitoring using a single repurposed imaging phased array [31], [32]. The use of a multi-element array enables a versatile, ultra-short focused transmit sequence with rapid electronic beam steering, as well as cavitation localization through power cavitation imaging (PCI). ThUS-mediated BBBO (ThUS-BBBO) has been studied in mice, where the rapid alternating steering angles (RASTA) pulse sequence was used to simultaneously induce bilateral BBB opening and enhance non-invasive dextran and adeno-associated virus (AAV) delivery to the brain through systemic injection [32]. In the same study, it was also shown that ThUS-BBBO can reversibly disrupt the brain immune homeostasis in a pulse-length dependent manner, suggesting that the method can be used as a neuro-immunotherapeutic, similar to conventional FUS-BBBO [13]. Minimal and reversible erythrocyte extravasation was reported, which was dependent on the pulse length of the applied ThUS sequence, with longer pulses exhibiting more hemorrhage at the 24-hour time point. This observation alludes to a potentially important role of pulse length in the behavior of microbubbles in ThUS-BBBO, and the associated effectiveness and safety of the resulting opening.

Notably, pulse length is not a factor in the calculation of the MI, therefore relying on the MI as the main predictor of safety and efficacy of FUS-BBBO may not be sufficient, particularly in the ultra-short pulse length regime. Previous studies have shown that successful BBB opening can be achieved with both long and short pulses, but short-pulse sequences require higher pressures to induce BBB opening volumes comparable to the long pulse ones [33], [34]. Our study with ultra-short ThUS pulses [32] demonstrated similar findings, where effective BBB opening and dextran or AAV delivery was achieved with a pressure approximately 1.0 MPa, corresponding to a MI of approximately 0.82. Unveiling the behavior of microbubbles under short pulses can therefore provide insight on the mechanism by which ThUS induces BBB opening.

To date, an extensive investigation into the nature of MB cavitation under ultra-short ThUS pulses has not been conducted. Distinguishing stable from inertial cavitation in the short-pulse regime has proven particularly challenging due to the extensive spectral leakage, or the spreading of energy associated with a particular frequency over multiple frequency bins which inherently occurs upon applying the Discrete Fourier Transform (DFT) on a time-limited or windowed signal [35]. As a result, harmonic and ultraharmonic microbubble responses, which appear as sharp peaks in long-pulse FUS, spread over a broader range of frequencies [36]. Therefore, the calculated cavitation dose from ThUS data may present a bias towards inertial cavitation due to increased calculated broadband activity, which, however, may not be directly linked to the damaging microbubble collapse associated with inertial cavitation. In order to fully understand the mechanisms of ThUS-BBBO drug delivery it is essential to characterize the nature of MB activity under ultra-short ThUS pulses.

In this study, we aim to characterize the cavitation activity under ultra-short ThUS pulses and the corresponding implications for *in vivo* AAV delivery. A phantom study was first carried out to understand the frequency composition of microbubble cavitation under ThUS and examine the extent of microbubble collapse under these ultra-short pulses [37]. The cavitation activity with ThUS-BBBO in an *in vivo* wild-type mouse model was also examined to understand the role of MBs on the BBB opening and the associated delivery of AAVs to the brain. The effect of microbubble size distribution, as well as the burst and pulse length on the resulting cavitation dose, BBBO volume and AAV delivery is also studied [38]. Overall, this study aims to investigate the types of microbubble cavitation involved in ThUS-BBBO and their associated bioeffects.

## Results

### ThUS pulses induce harmonic and broadband cavitation

In the phantom studies, the cavitation dose was calculated for baseline and microbubble conditions from the frequency domain data obtained using a PCD as well the received data from the P4-1 array. Qualitative comparison between baseline and microbubble measurements shows increased signal amplitude at higher frequencies for all pulse lengths when microbubbles are introduced to the flow channel (Fig. **2A-D**). In particular, for the 10– and 20-cycle conditions, this signal amplitude rise is centered around the harmonic frequencies whereas for the shorter pulse length measurements there is significantly more spectral broadening. Additionally, the significant spectral leakage observed around the fundamental frequency for the 1.5-cycle conditions led to calculation of higher ultraharmonic and broadband cavitation doses (Fig. **2E-F**). Quantitatively, the contribution of harmonic frequencies to the total cavitation dose reaches 66.5% for 20 cycles, which is approximately 4.6 times higher than the 1.5 cycle conditions.

**Figure 1.**
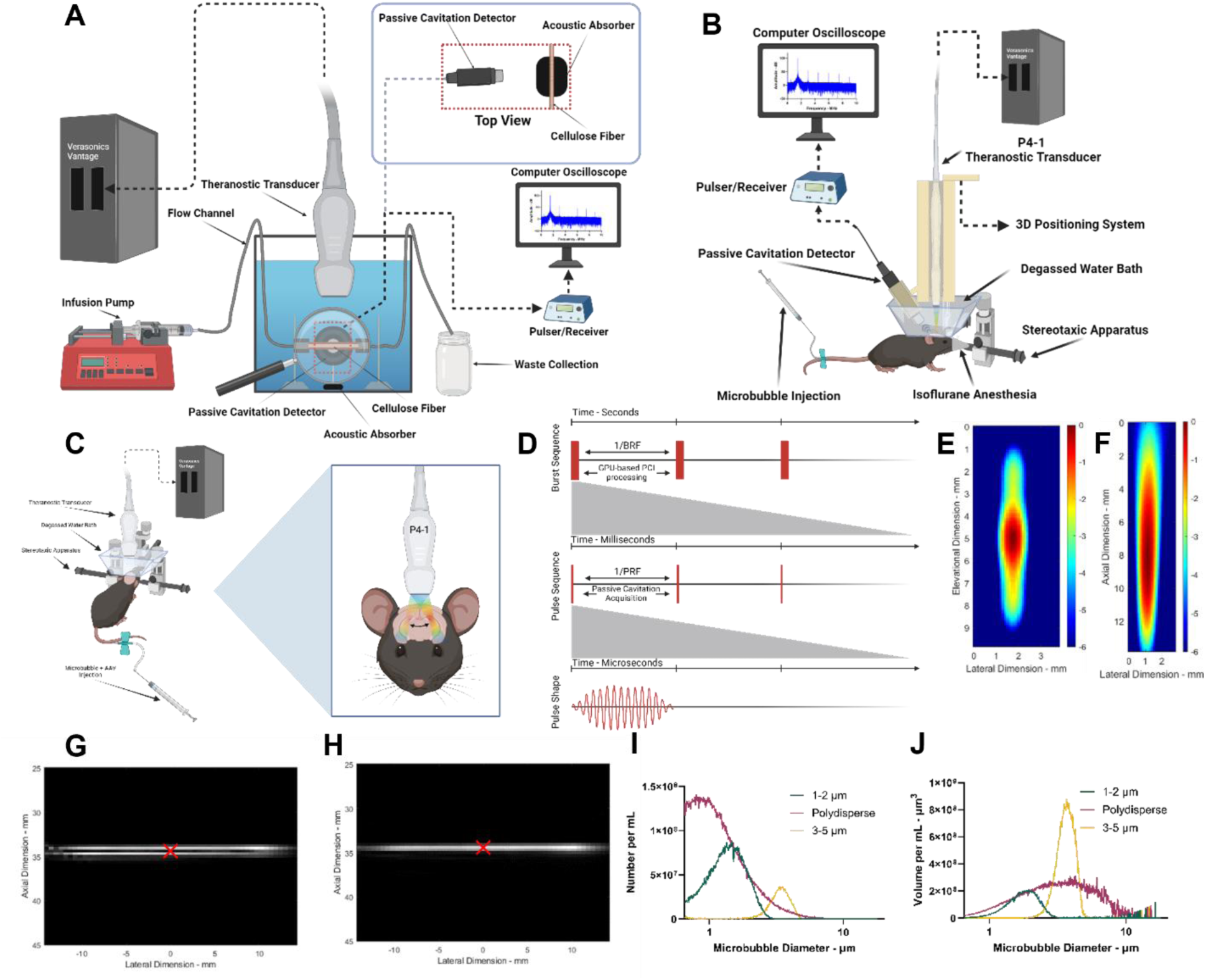
**A**) Schematic of the flow phantom setup for quantification of ThUS cavitation with a passive cavitation detector. **B)** Schematic of the *in vivo* ThUS-BBBO mouse study; in addition to monitoring via power cavitation imaging, cavitation was quantified using a passive cavitation detector. **C)** Schematic of the *in vivo* ThUS-BBBO and AAV delivery studies, where the RASTA sequence was used for simultaneous bilateral BBBO with 1.5 cycles on the left hemisphere and 10 cycles on the right hemisphere. **D)** General schematic of ThUS pulse sequences showing the composition of bursts and pulses. **E-F)** Pressure field map of the P4-1 focused transmits at 35 mm focusing distance on the lateral-elevational plane (E) and on the lateral-axial plane (F). **G-H)** B-mode image of the flow tube with deionized water (G), or microbubble solution (H) used for targeting of the ThUS focus on the cellulose flow channel. **I-J)** Microbubble distribution of the in-house synthesized lipid-shelled microbubbles expressed as number per mL (I) or volume per mL (J).

**Figure 2.**
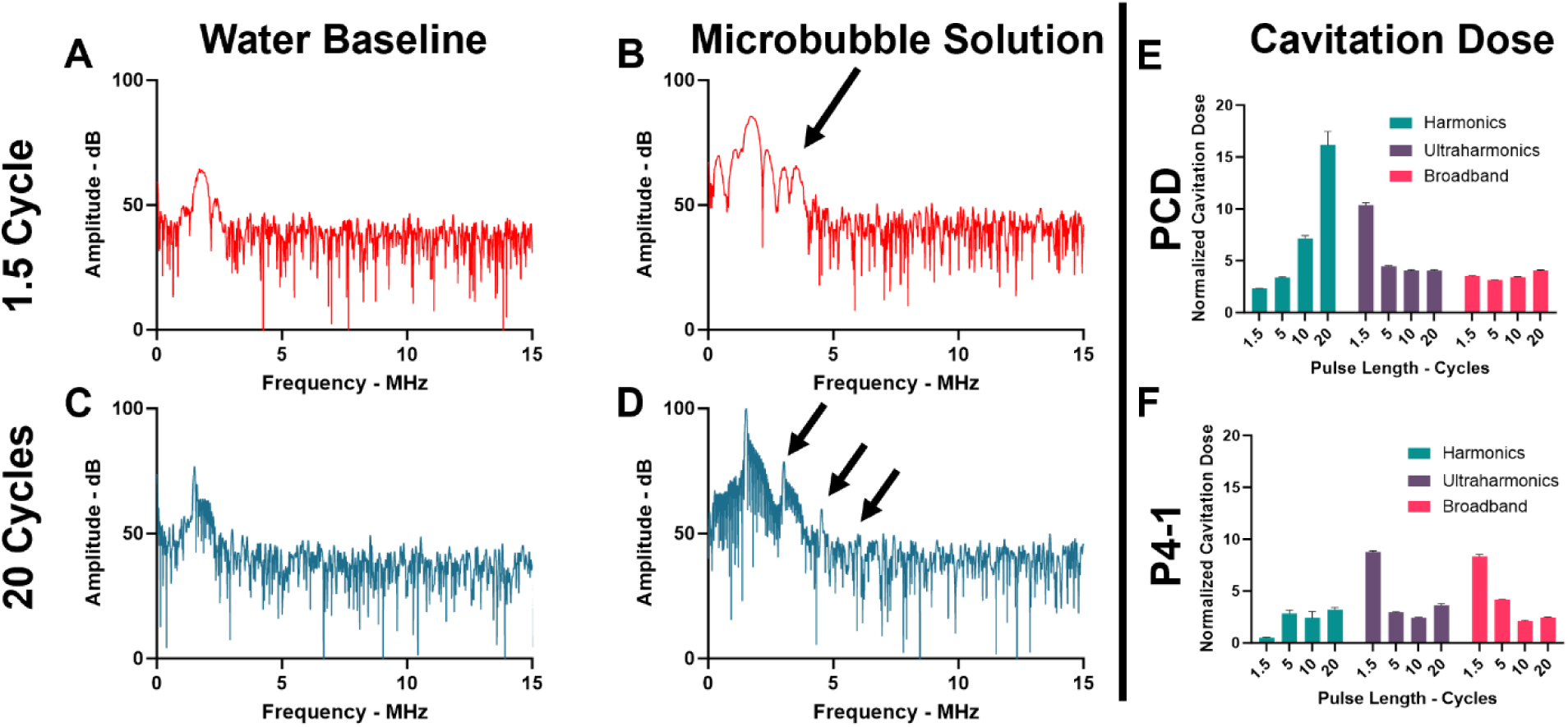
Microbubble behavior under ThUS in free field, detected by PCD in the flow phantom. Measurements were obtained by injecting degassed, deionized water or microbubble solution through the phantom at a flow rate of 100 μL/min. **A-B)** Frequency domain data for the first 1.5-cycle pulse of a burst with deionized water (A) or microbubble solution (B) flow in the phantom. Increased amplitude at higher frequencies (black arrow on (B)) is observed when microbubbles are introduced, however this signal is not centered around harmonic frequencies due to significant spectral leakage. **C-D)** Frequency domain data for the first 20-cycle pulse of a burst with deionized water (C) or microbubble solution (D) flow in the phantom. Increases in higher frequency signal are centered around the harmonic frequencies as indicated by black arrows on (D). **E-F)** Normalized cavitation dose calculated for all pulse lengths from the PCD (E) and P4-1 (F) data, showing increases in harmonic cavitation dose with pulse length, and decreases in ultraharmonic and broadband cavitation doses. The cavitation dose from P4-1 data does not capture harmonic frequencies as effectively as the PCD and is more heavily skewed towards ultraharmonic and broadband cavitation.

Comparison between the cavitation dose calculated from the PCD and the P4-1 data shows a significant dominance of broadband and ultraharmonic cavitation in the P4-1 signal, particularly for the shorter pulse lengths (Fig. **2F**). Additionally, the harmonic components of the P4-1 signal are increasing with pulse length, albeit to a much smaller extent compared to the PCD data (Fig. **2E-F**). The P4-1 signal is therefore mostly made of ultraharmonic and broadband components, and there is not a good representation of harmonic components.

### Microbubble collapse depends on ThUS pulse length

To determine whether the significant dominance of broadband cavitation dose is due to microbubble collapse or merely a consequence of the DFT-induced spectral leakage, the data obtained by PCD were analyzed in the time domain. The change in the time-domain signal amplitude over an entire ThUS burst was observed under water or microbubble solution flow at a rate of 100 μL/min. For baseline measurements with water, the amplitude remained largely unchanged throughout the duration of the burst (Fig. **3A**); however, for microbubble measurements the signal strength was not only much larger than baseline but showed an exponential-like decay over the 100 ThUS pulses (Fig. **3B-C**). Furthermore, the rate of decay, as well as the final signal amplitude, depended on pulse length (Fig. **3G**, Tab. **2**). The shortest, 1.5-cycle pulse exhibited the smallest amplitude drop (Fig. **3C**), whereas the 20-cycle shows a sharper and more drastic signal decrease (Fig. **3B**). B-mode images before and after the application of ThUS pulses also show the differences in microbubble collapse at different pulse lengths (Fig. **4**).

**Figure 3.**
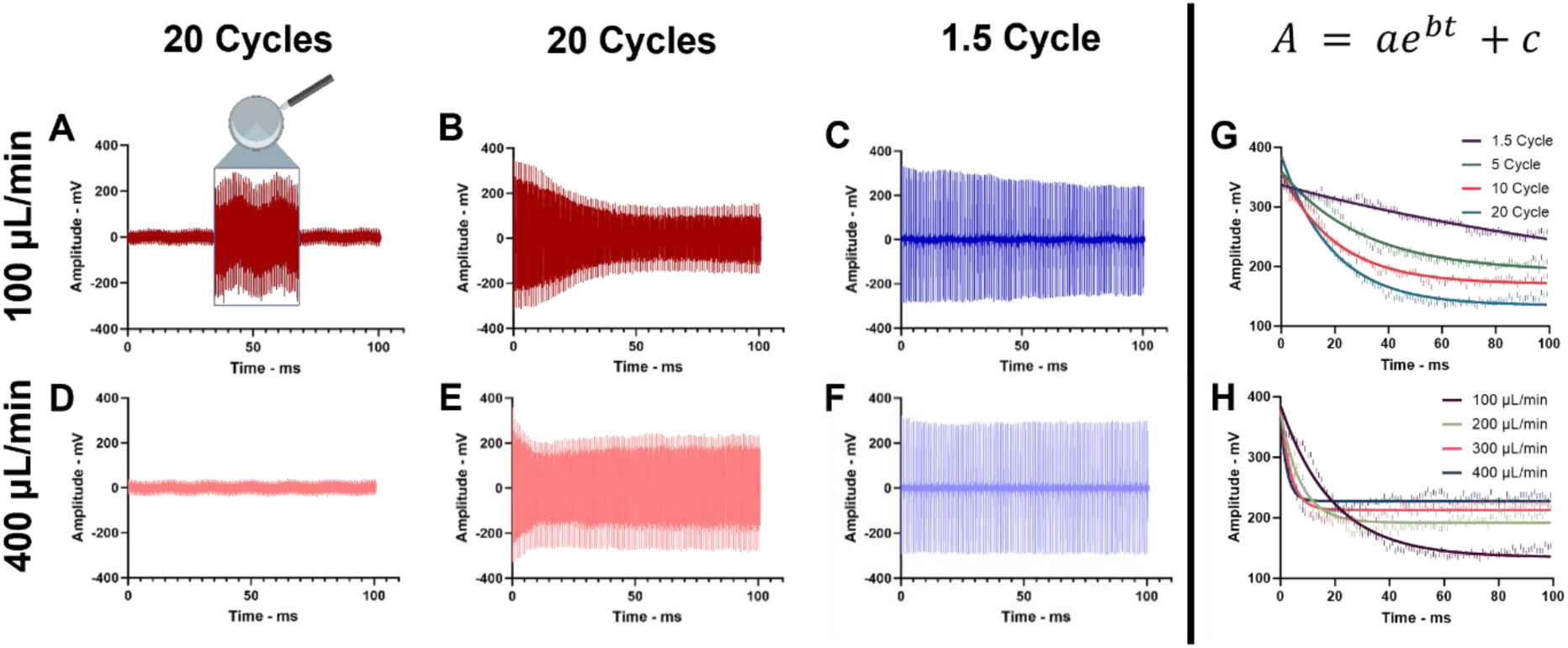
PCD signal during a ThUS burst in the time domain for different pulse lengths and flow rates. **A-B)** Time domain PCD signal from the flow phantom under baseline conditions (A) or microbubble infusion at 100 μL/min (B) and sonication with 20-cycle pulses. Baseline signal (A) shows minimal fluctuation due to radio frequency noise, while microbubble signal (B) decreases exponentially over the course of the burst. **C)** Time domain PCD signal from the flow phantom with microbubble infusion at 100 μL/min and 1.5-cycle pulses. Signal decrease over the course of the burst is observed, albeit being smaller compared to the 20-cycle case. **D-E)** Time domain PCD signal from the flow phantom under baseline conditions (D) or microbubble infusion at 400 μL/min (E) and sonication with 20-cycle pulses. Baseline signal (D) again shows minimal fluctuation due to radio frequency noise, however microbubble signal (E) decreases to a smaller extent compared to the same pulse length sonication with slower flow rate. **F)** Time domain PCD signal from the flow phantom with microbubble infusion at 400 μL/min and 1.5-cycle pulses shows minimal change in signal amplitude over the course of the burst. **G)** Exponential fit of the time domain peak-positive signal amplitude for different pulse lengths when microbubbles were infused at 100 μL/min. Increased pulse length resulted in increased rate of signal amplitude decrease, and lower final amplitude values. **H)** Exponential fit of the time domain peak-positive signal amplitude for different flow rates when microbubbles were infused, and 20-cycle pulses were applied. Increased flow rates resulted in higher final amplitude values. Data points in **G)** and **H)** were fitted with the exponential model A = ae^bt^+c on MATLAB.

**Figure 4.**
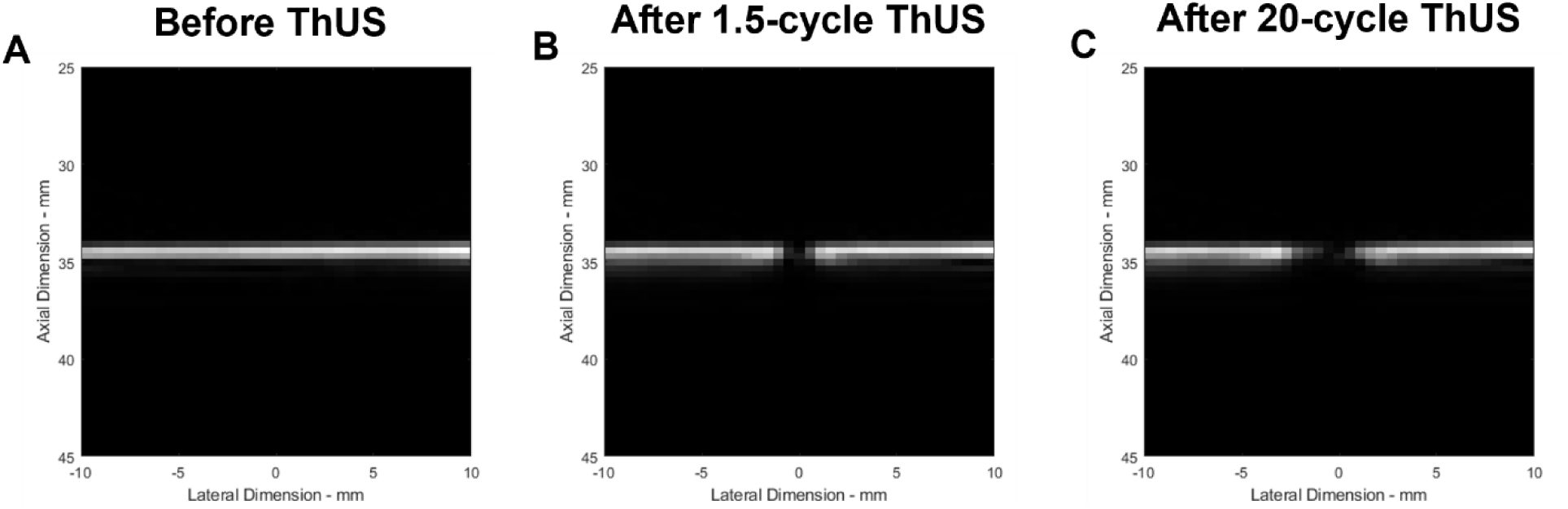
**A-C**) B-mode image at 2.5 MHz before ThUS with stationary microbubbles in the flow channel before ThUS (A), or after 5 bursts with 1.5-cycle (B) or 20-cycle (C) focused transmits. Signal loss in the center of the channel indicates microbubble depletion due to ThUS. The signal loss in the center of the channel is more extensive for the 20-cycle case compared to 1.5 cycle, indicating increased microbubble collapse.

Next, we sought to understand the effect of the flow rate on the signal amplitude decay. Figure **3H** shows that the decay rate, as well as the extent of signal amplitude decrease are affected by the microbubble flow rate. An exponential decay fit was applied to quantify the extent of microbubble collapse for the various pulse length and flow rate conditions. Table **2** summarizes the exponential fit parameters for the different experiments. It is indeed observed that the faster flow rates result in a lower overall decrease in the cavitation signal amplitude over the 100 focused pulses of a burst. Interestingly, the increase in flow rate also results in an increase in the rate of the exponential-like decay of the time-domain signal.

### ThUS induces harmonic and broadband cavitation *in vivo* in mice

Analysis of the PCD data from the first *in vivo* mouse study revealed the components of the microbubble signal following ThUS-BBBO. Upon injection of the microbubbles, a sharp increase in the strength of the harmonic components of the PCD signal was observed for both 1.5 cycle and 10 cycle conditions, which were maintained throughout the sonication procedure (Fig. **5D-E**). This increase in harmonic cavitation dose was larger for the 10-cycle pulse length compared to the 1.5-cycle, up to 24% in the former compared to 5.5% in the latter. In addition to this, an increase in the broadband cavitation dose was also observed for both pulse lengths, albeit to a smaller extent compared to the harmonic dose; up to 3% for the 10-cycle conditions and 2.5% for the 1.5-cycle conditions. Interestingly, ultraharmonic cavitation dose shows a decrease when the microbubbles are injected, at both pulse lengths. We attribute this observation to the overall weak amplitude of ultraharmonic peaks and to the increased spectral leakage due to the short pulse length, which makes ultraharmonic cavitation difficult to resolve from broadband signal.

**Figure 5.**
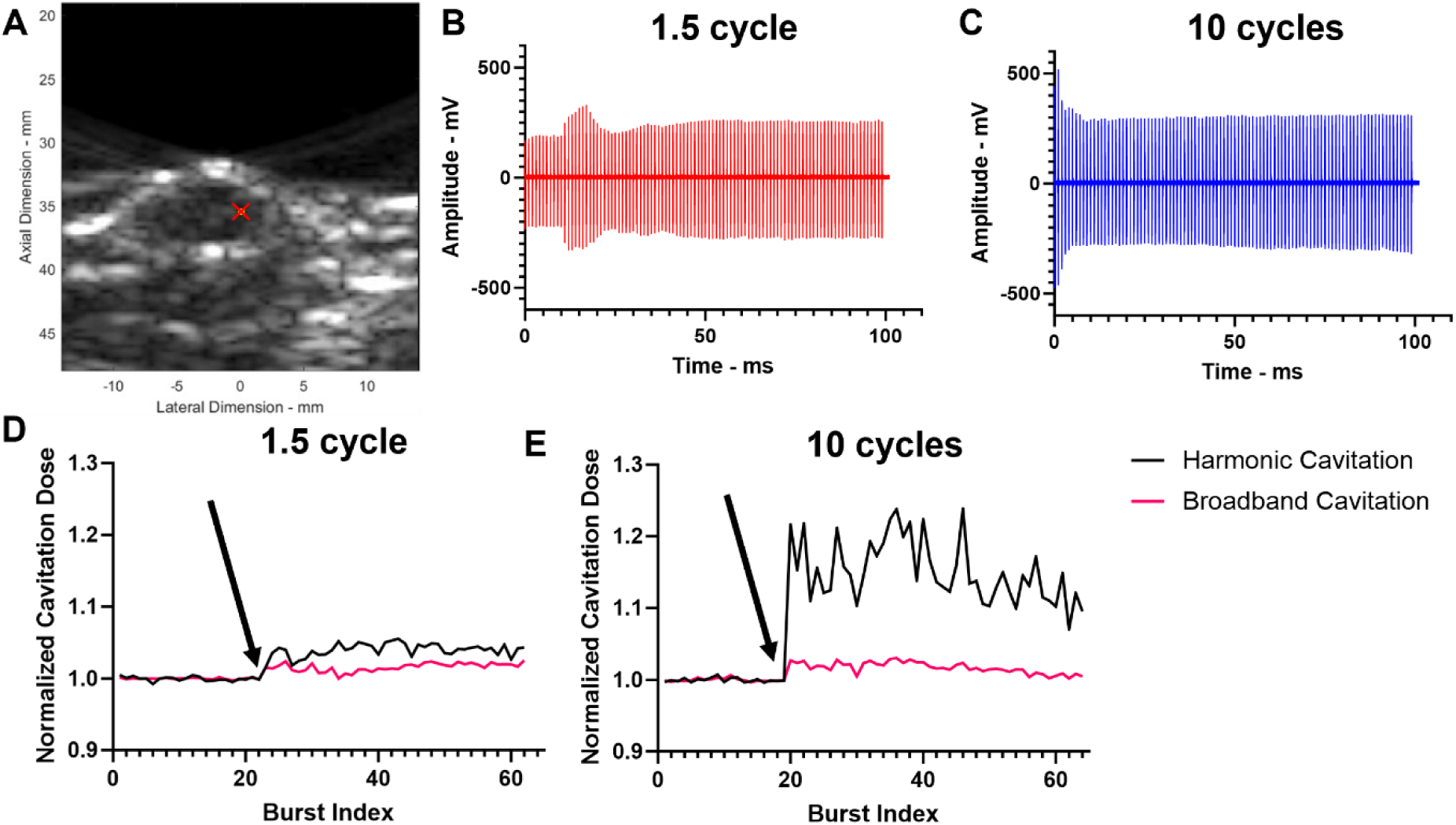
ThUS cavitation measurements from *in vivo* mouse study with PCD. **A)** B-mode imaging at 2.5 MHz was used to target the ThUS focus over the hippocampus of C57BL/6 mice. The red cross indicates the center of the ThUS focused beam. **B)** Time domain signal of a burst received by PCD after microbubble injection and sonication 1.5-cycle (B) or 10-cycle (C) ThUS. The cavitation signal peaks to a larger amplitude for the 10-cycle case and then decreases faster for the remainder of the burst. **D-E)** Normalized cavitation dose during the entire sonication with 1.5 cycle (D) or 10 cycles (E). Black arrow indicates the timepoint where microbubbles were injected. A significant increase in harmonic cavitation dose is observed for the 10-cycle case compared to 1.5 cycle, while broadband cavitation also increases, albeit to a smaller extent.

Time domain signal analysis showed a similar microbubble behavior as the phantom experiments. For the shorter, 1.5 cycle pulse lengths, the drop in signal amplitude is smaller, and recovers faster than the 10-cycle case (Fig. **5B**). The longer pulse length exhibits an exponential-like signal decrease over the 100 pulses, similar to the phantom experiments, indicating a faster rate of microbubble collapse than the 1.5 cycle (Fig. **5C**).

### ThUS cavitation, BBBO volume, and AAV delivery increase with microbubble size

The following mouse studies aimed to further characterize the relationship of ThUS cavitation to the BBB opening and resulting AAV delivery. In these studies, cavitation dose was calculated by summation of the pixels in PCI maps. Firstly, the effect of microbubble size distribution on the resulting ThUS cavitation dose, BBBO volume and AAV delivery was investigated. Three groups of mice received one of three microbubble formulations co-injected with the AAV9-CAG-GFP construct. The resulting BBB opening was evaluated using gadolinium contrast-enhanced T1-weighted MRI (Fig. **6A-C**). The average BBB opening volumes for the 1-2 μm, polydisperse, and 3-5 μm microbubble groups were 10.17 mm^3^, 93.45 mm^3^, and 158.34 mm^3^ respectively. Larger BBB opening volumes were observed on the right hemisphere, which was sonicated with 10 cycle pulses, as opposed to 1.5 cycle on the left hemisphere. It was also observed that larger microbubbles induce a higher cavitation dose, as calculated from the PCI maps (Fig. **6G-I**). The BBB opening volume shows a positive linear relationship with baseline-normalized PCI cavitation dose (R^2^=0.60, p=0.0007, Fig. **6M**), as well as with total PCI-cavitation dose (R^2^=0.70, p<0.001, Fig. **6N**) across the three microbubble groups, while the relationship between baseline-normalized cavitation and BBB opening volume within groups only was significant for the polydisperse microbubble group (R^2^=0.60, p=0.0007, Fig. **6K**). Moreover, no significant correlations within or between groups was found between the frequency domain-derived harmonic, ultraharmonic, and broadband cavitation doses and BBB opening volume (Fig. **S1A-C**).

**Figure 6.**
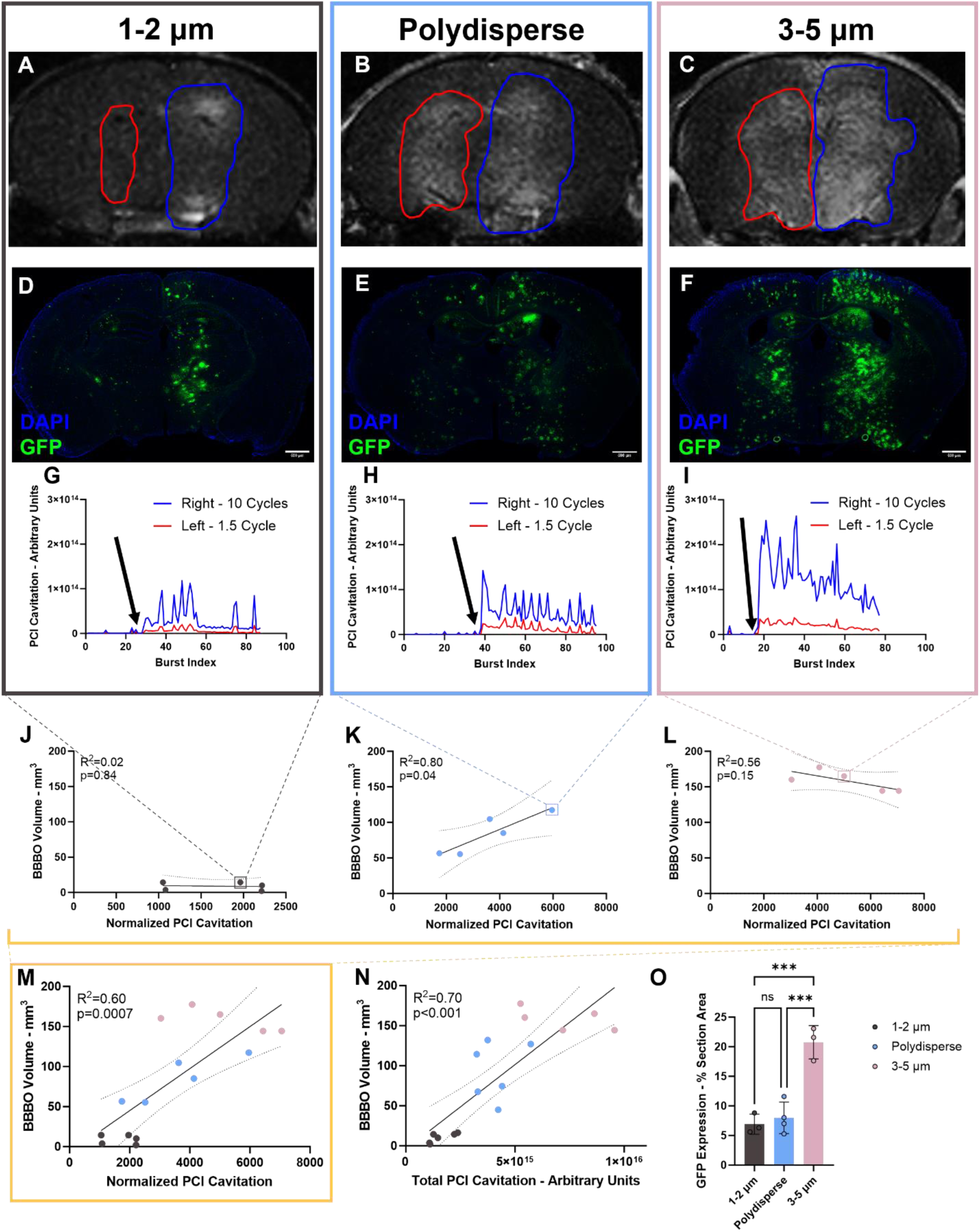
Results of the ThUS-mediated AAV delivery study with different microbubble size distributions. **A-C)** Contrast-enhanced T1-weighted MRI for a mouse injected with 1-2 μm (A), polydisperse (B), and 3-5 μm (C) MBs and sonicated with 1.5 cycle on the left and 10 cycles on the right. BBB opening is indicated by red (1.5 cycle) and blue (10 cycle) contours. The contrast enhancement region is larger and with higher intensity as microbubble size increases. **D-F)** Fluorescence microscopy of a representative section from a mouse injected with 1-2 μm (D), polydisperse (E), and 3-5 μm (F) MBs and sonicated with 1.5 cycle on the left and 10 cycles on the right. DAPI is stained in blue and GFP in green. GFP expression is higher on the right side and increases with larger MB size. **G-I)** PCI-derived cavitation dose over the course of the sonication for a mouse injected with 1-2 μm (G), polydisperse (H), and 3-5 μm (I) MBs and sonicated with 1.5 cycle on the left and 10 cycles on the right. Blue and red lines represent the cavitation dose calculated from 10– and 1.5-cycle pulses respectively. Black arrows indicate the timepoint where MBs were injected. Cavitation dose is higher for 10-cycle pulses and increases with MB size. **J-L)** Correlation of BBB opening volume with normalized PCI cavitation dose within each group of mice injected with 1-2 μm (J), polydisperse (K), and 3-5 μm (L) MBs. Linear regression analysis was performed, with a significant correlation only being noted for the polydisperse MB case (K). **M-N)** Correlation of BBB opening volume with baseline-normalized (M) and total (N) PCI-derived cavitation dose across the three MB groups. Linear regression analysis indicates a significant correlation across groups for both normalized and total cavitation dose. (M) is a collection of the plots in (J-L). **O)** GFP expression on brain sections of mice from the MB study, expressed as a percentage of the total section area. One-way ANOVA with post-hoc Tukey’s multiple comparisons test was performed indicating statistically significant increase in GPF expression for the 3-5 μm MB group compared to the other two groups. Statistical significance across all figures was set for p<0.05; ***:p<0.001.

Furthermore, the larger microbubble groups resulted in increased transgene expression in the brain. The GFP coverage in each section was quantified using a MATLAB script and the maximum percentage coverage per section was found to be 6.92 ± 0.98 %, 7.98 ± 1.33 %, and 20.76 ± 1.62 % for the 1-2 μm, polydisperse, and 3-5 μm microbubble group respectively (Fig. **6O**). Like BBB opening, the transgene expression was larger on the right hemisphere which was sonicated with 10-cycle pulses (Fig. **6D-F**).

Finally, safety was assessed for each microbubble group using T2*-weighted MRI imaging, as well as H&E staining. T2*-weighted images revealed a small region of hypo-intensity representing mild hemorrhage on the right hemisphere of mice injected polydisperse and 3-5 μm microbubbles without any significant findings for the 1-2 μm group (Fig. **S2A-C**). H&E staining also revealed mild erythrocyte extravasation for all microbubble groups on the right hemisphere, and smaller areas of microhemorrhage on the left hemisphere of mice injected with polydisperse and 3-5 μm bubbles (Fig. **S2D-L**). Despite this finding, the mice did not show any behavioral deficiencies and survived for the entire three-week survival period.

### ThUS burst length affects cavitation dose and AAV delivery

Next, the effect of burst length and burst repetition frequency on the resulting BBB opening, cavitation dose and AAV delivery was investigated. The number of pulses per burst, i.e. the burst length, and burst repetition frequency for each group were adjusted to an effective duty cycle of 50 pulses per second, so that each group receives approximately the same number of pulses throughout the 2-minute sonication. The median number of pulses throughout the sonication was 5675, 5650, and 6400 for the 50, 100, and 200 pulse-per-burst groups respectively. Despite the larger number of pulses applied with the longer burst length group, the BBB opening volumes were the smallest, averaging 55.25 mm^3^, compared to 93.45 mm^3^, and 152.6 mm^3^ for the 100 and 50 pulse-per-burst groups respectively.

The cavitation dose was subsequently calculated for each mouse from the PCI maps. The BBB opening volume correlated negatively with total PCI-derived cavitation dose across groups (R^2^=0.48, p=0.003, Fig. **7N**), as the 200-pulse group had the highest total cavitation dose due to the largest number of pulses, but the lowest BBB opening volumes. Conversely, when the PCI cavitation dose is normalized by the baseline, i.e. the pulses before microbubble injection, there is a positive correlation across groups between cavitation dose and BBB opening volume (R^2^=0.70, p<0.0001, Fig. **7M**). For normalized PCI-derived cavitation dose, a significant positive correlation was found within the 100-pulse group (R^2^=0.60, p=0.0007), similar to what was shown in our previous study [32], but no other groups showed significant correlations. Positive correlations across groups were found between the frequency domain-derived harmonic, ultraharmonic and broadband cavitation doses and BBB opening volumes, without however any significant correlation within any of the groups (Fig. **S1D-F**).

**Figure 7.**
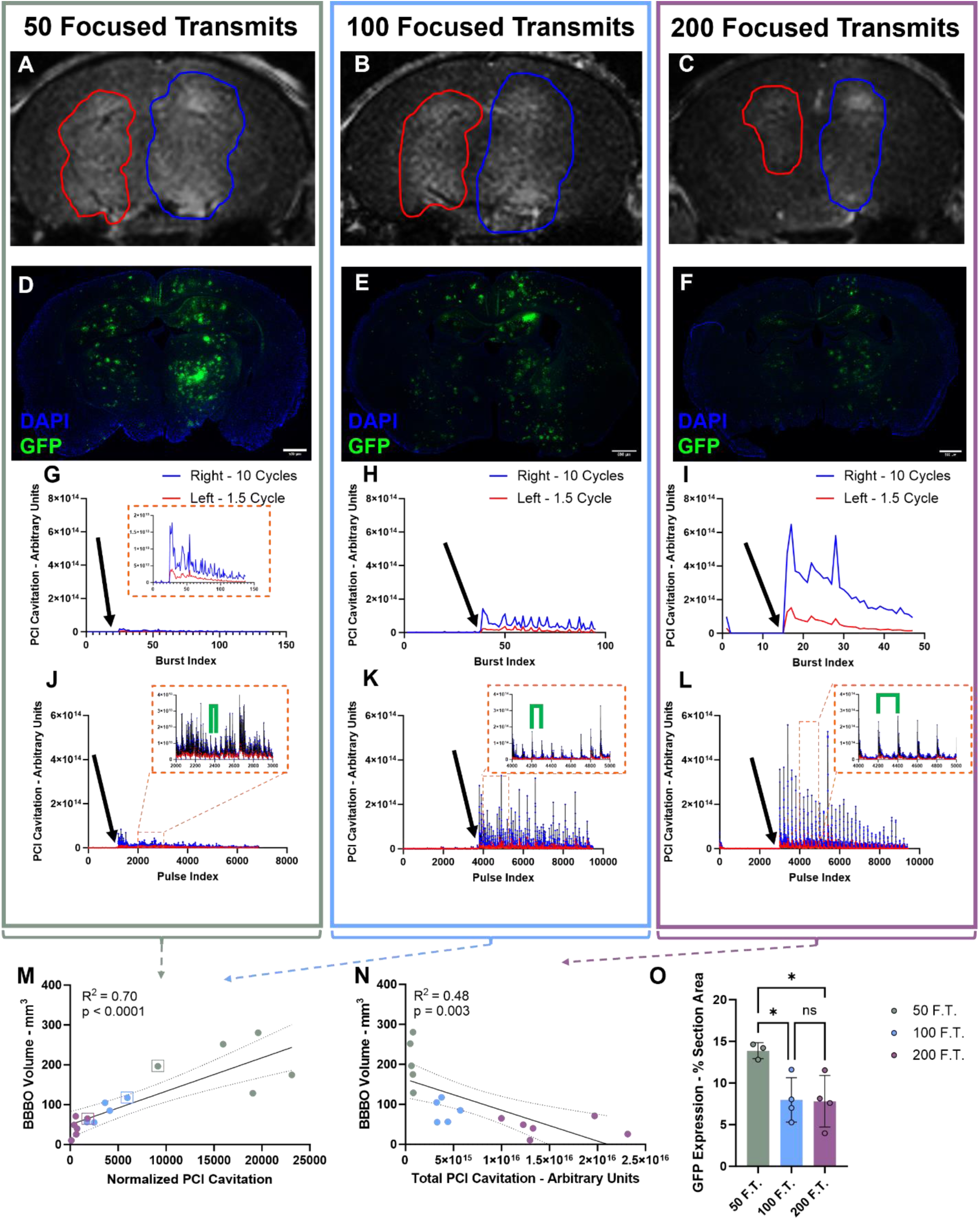
Results of the ThUS-mediated AAV delivery study with different burst sequences. **A-C)** Contrast-enhanced T1-weighted MRI for a mouse sonicated 1.5 cycle on the left and 10 cycles on the right and 50 (A), 100 (B), and 200 (C) focused transmits per burst. BBB opening is indicated by red (1.5 cycle) and blue (10 cycle) contours. The contrast enhancement region is larger and with higher intensity for shorter burst lengths. **D-F)** Fluorescence microscopy of a representative section from a mouse sonicated with 1.5 cycle on the left and 10 cycles on the right and 50 (D), 100 (E), and 200 (F) focused transmits per burst. DAPI is stained in blue and GFP in green. GFP expression is consistently higher on the right side and increases with shorter burst length. **G-I)** PCI-derived cavitation dose over the course of the sonication for a mouse sonicated 50 (G), 100 (H), and 200 (I) focused transmits per burst. Blue and red lines represent the cavitation dose calculated from 10– and 1.5-cycle pulses respectively. Black arrows indicate the timepoint where MBs were injected. Inset in (G) shows the plot in (G) zoomed 10 times in the vertical axis. Cavitation dose is consistently higher for 10-cycle pulses and increases with burst length. **J-L)** PCI-derived cavitation dose for each individual pulse throughout the sonication for 50 (J), 100 (K), and 200 (L) focused transmits per burst. Blue and red points indicate 10– and 1.5-cycle pulses respectively. Black arrows indicate the timepoint where MBs were injected. Insets in (J-L) show the respective plots zoomed over 1000 pulses in the horizontal axis. In all plots, cavitation dose peaks at the beginning of every burst and subsequently decreases to near-baseline levels. The green brackets in the insets indicate the distance between consecutive cavitation peaks, showing that for 50-pulse bursts, there are more frequent cavitation peaks compared to the larger burst lengths. **M-N)** Correlation of BBB opening volume with baseline-normalized (M) and total (N) PCI-derived cavitation dose across the three bursts sequence groups. Linear regression analysis indicates a significant correlation across groups for both normalized and total cavitation dose, albeit a positive correlation for normalized PCI cavitation (M) and a negative correlation for total cavitation (N) **O)** GFP expression on brain sections of mice from the burst sequence study, expressed as a percentage of the total section area. One-way ANOVA with post-hoc Tukey’s multiple comparisons test was performed indicating statistically significant increase in GPF expression for 50-pulse-per-burst group compared to the other two groups. Statistical significance across all figures was set for p<0.05; *:p<0.05.

The AAV transgene expression 3 weeks after sonication was also assessed. Similar to BBB opening volume, GFP expression was largest for the 50-pulse group, reaching maximum section coverage of 13.88 ± 0.55 % compared to 7.98 ± 1.33 % and 7.81 ± 1.55 % for the 100– and 200-pulse groups respectively (Fig. **7O**).

Finally, T2*-weighted MRI from the day of sonication and H&E-stained sections of brains collected 1 day after sonication were used to assess the safety of each burst sequence. Regions of hypointensity, indicating hemorrhage were observed on T2* images of mice sonicated with 50 and 100 focused transmits per pulse (Fig. **S3A-B**), while small areas of erythrocyte extravasation were present on the H&E-stained sections for all conditions (Fig. **S3D-L**). Similar to the microbubble study, the mice survived for the entire three-week survival period, despite the noted microhemorrhage.

### AAV serotype and injection dose affect cell types and number of cells transduced

Finally, we studied the effect of AAV serotype on the types of cells transduced, as well as the overall transgene expression efficiency following ThUS-BBBO. All AAV constructs contained the same, ubiquitous CAG promoter, and encoded the GFP protein. It was observed that AAV9 yielded significantly larger transgene expression compared to both AAV5 and AAV2 for the same injected dose of 3 x 10^11^ gc/mouse, with maximum section coverage of 16.96 %, 2.01 %, and 1.11 % for AAV9, AAV2 and AAV5 respectively (Fig. **8E**). Additionally, the latter two serotypes were better at specifically transducing neurons, whereas with AAV9 different cell types such as neurons and astrocytes expressed the GFP transgene.

**Figure 8.**
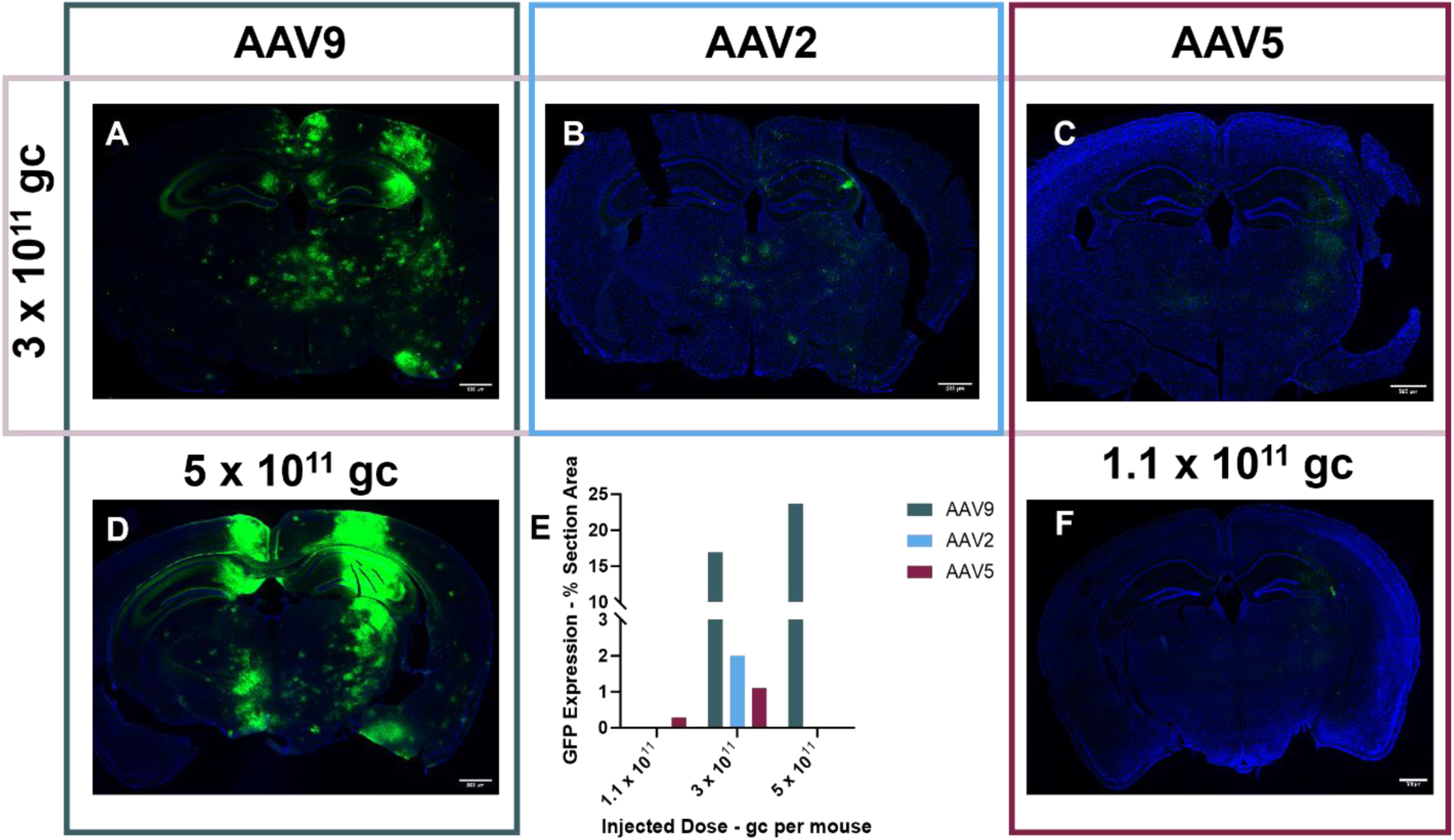
Results of the ThUS-mediated AAV delivery study with different AAV serotypes and injection doses. **A-C)** Fluorescence microscopy of a representative brain section from mice injected with systemic dose of 3 x 10^11^ gc/mouse and the AAV9 (A), AAV2 (B), and AAV5 (C) serotype, all encoding GFP with the CAG promoter. GFP expression is significantly higher with AAV9 compared to the other two serotypes. **D)** Fluorescence microscopy of a brain section from a mouse injected with 5 x 10^11^ gc of AAV9-CAG-GFP. The increased fluorescence intensity in the targeted regions compared to (A) indicates increased transduction of the reporter gene at the higher systemic dose. **E)** GFP expression on brain sections of mice injected with different serotypes and doses. AAV9 shows significantly higher expression compared to the other two AAVs (Note the discontinuous vertical axis). n=1 per group. **F)** Fluorescence microscopy of a brain section from a mouse injected with 1.1 x 10^11^ gc of AAV5-CAG-GFP, showing limited GFP expression in neurons within the targeted region, and lower expression compared the higher dose in (C).

Furthermore, increasing the dose of AAV resulted in increased GFP expression in the areas of BBB opening. For AAV9, the transgene expression increased from 16.96 % to 23.68 % for injection doses of 3 x 10^11^ gc/mouse and 5 x 10^11^ gc/mouse respectively, whereas for AAV5, the GFP expression increased from 0.29 % to 1.11 % between 1.1 x 10^11^ gc/mouse and 3 x 10^11^ gc/mouse (Fig. **8E**).

Notably, increasing the dose of AAV did not seem to have an effect on the health of the mice throughout the 3 weeks that they were allowed to survive. However, mice injected with AAV9 showed significant peripheral organ transduction compared to other serotypes, particularly in the liver. Figure **S4** shows the difference in liver transduction between the mice injected with different serotypes.

## Discussion

This is the first comprehensive study on the characterization of MB activity under ultra-short ThUS pulses used in a flow phantom and *in vivo*. While most prior studies categorize microbubble activity as stable or inertial based on the harmonic/ultraharmonic or broadband frequency response, it was demonstrated here that this classification is neither appropriate nor sufficient in the short-pulse ThUS regime. Due to significant spectral leakage in the frequency domain, ultra-short pulse lengths of 1.5 cycle exhibit increased broadband cavitation dose, which is typically associated with microbubble collapse and microjetting. However, analysis of signal amplitude over time reveals that despite the increased broadband signal in the frequency domain, there is not a significant decrease in the signal strength for the shortest pulse length. On the other hand, longer pulse lengths such as 20 cycles, which show significantly higher harmonic cavitation in the frequency domain, show a drastic decrease in signal amplitude over time, which is evidence of increased transient cavitation. These results serve as evidence that for the ultra-short pulse length regimes, the characterization of inertial cavitation and microbubble collapse from broadband signal is not a sufficient metric.

This signal amplitude decrease in the time domain is attributed mainly to microbubble collapse for the following reasons: Firstly, a variation in signal is not observed in experiments with identical flow conditions but an absence of microbubbles, showing that this effect is due to the backscattered and secondary microbubble signal. Moreover, and particularly for the flow phantom experiments with 100 μL/min flow rate, the rate of microbubble infusion is significantly slower than the pulse repetition frequency and the ultra-short pulse length, indicating that the quantity of microbubbles under the ThUS focus does not get replenished over a single burst. Finally, since there is continuous flow and due to the small diameter of the channel, the microbubbles are restricted within the focal volume, thus minimizing the effects of acoustic radiation forces and the mere displacement of microbubbles away from the focus. Therefore, the decrease in time-domain signal amplitude is interpreted as microbubble collapse.

An interesting observation from the phantom and *in vivo* studies is that the extent of microbubble collapse, as evidenced by the decrease in signal amplitude in the time domain, is dependent on both the pulse length and microbubble flow rate. It is observed that increasing the pulse length while keeping all other parameters constant leads to increased microbubble collapse as well as increased cavitation dose. This increased cavitation activity translates to a larger BBB opening volume *in vivo*, as well as more extensive microhemorrhage, which is consistent with our previous study [39]. This result points to a dependence of the BBB opening volume and safety on the total energy deposited to the tissue from cavitating microbubbles, rather than simply on the MI. It also demonstrates that the mechanism of ThUS-mediated BBB opening is primarily transient cavitation driven, as BBB opening occurs following microbubble depletion after few ThUS pulses, particularly for the longer 10-cycle pulse length. This transient cavitation may be due to the high MI required for BBB opening with ultra-short pulses, and the effect may be exacerbated with increased pulse length as microbubbles are being exposed to high MI conditions for longer, leading to more probability of collapse. In our *in vivo* studies, mice received the same MI on both hemispheres, but the hemisphere treated with 10 cycles showed microhemorrhage, demonstrating that the increased duty cycle of ThUS contributed to the damaging effects, as well as the enhanced BBB opening and AAV delivery. It is also noteworthy that our previous study [32] has shown that the microhemorrhage observed after ThUS is transient and mostly resolves within 4 days. Therefore, for optimized AAV delivery, the pulse length can be adjusted to maximize BBB opening volume while staying below the damage threshold.

One of the main advantages of using a single theranostic transducer is the ability to create cavitation maps of high resolution through power cavitation imaging (PCI). Our published work has demonstrated superior PCI resolution with shorter pulse lengths [32], with better cavitation localization at 1.5 cycle compared to 10 cycles. However, we demonstrate here that spectral leakage decreases with increased pulse length, making the distinction of cavitation types more reliable at longer pulse lengths. There is therefore a tradeoff between high resolution microbubble localization and accurate cavitation characterization. Depending on the study design, an optimal pulse length can be chosen to prioritize PCI resolution or cavitation type characterization as well as optimize drug delivery efficiency and microhemorrhage.

In the flow phantom studies and the first *in vivo* study, cavitation was characterized both using the PCD and the P4-1 data. The use of a separate system for cavitation detection enables the acquisition of high frequency harmonics, which tend to be better indicators of microbubble activity. The PCD data served as the ground truth for cavitation dose calculation as they were obtained at a sufficiently high sampling frequency and the bandwidth of the PCD was sufficiently large to receive the higher frequency signals. The cavitation dose calculation from the P4-1 data could only include the 3 and 4.5 MHz harmonic frequencies, which may also include significant contributions from the skull and surrounding tissue. For this reason, the frequency-domain cavitation dose calculated from P4-1 data may not accurately represent microbubble activity, especially without the application of SVD filtering. Using the PCI maps as a proxy for cavitation dose calculation may offer a better measure of the microbubble activity, as the skull, and non-moving tissue contributions are being removed by the SVD filter. In our *in vivo* studies where a separate PCD was not used for cavitation detection, the PCI method of cavitation calculation proved more indicative of correlating with BBB opening volume compared to the frequency domain calculation, thus showing that PCI-derived cavitation dose can be a sufficient metric for BBBO volume and AAV delivery prediction. Therefore, in the microbubble and burst sequence *in vivo* studies with AAV delivery, the PCI-derived cavitation dose was primarily used as a cavitation metric.

Our flow phantom studies had shown that the majority of cavitation activity occurs at the beginning of a burst, as the microbubbles are depleted within the first few pulses. The burst length and BRF were therefore varied to allow for microbubble replenishment in the focal volume and subsequently increase the effects of cavitating microbubbles on the BBB. It was shown that irrespective of burst length, the majority of cavitation occurs during the first pulses of a burst, and decays in subsequent pulses (Fig **7J-L**). When the PCI-derived cavitation dose was normalized by the baseline, the group with the shortest burst length and fastest burst repetition frequency exhibited the highest cavitation doses, which correlated with the highest BBB opening volumes and AAV transgene expression. This observation therefore demonstrates the importance of having sufficient microbubbles in the vasculature to achieve increased BBB opening and permeability.

In the *in vivo* microbubble study, the ThUS-BBBO volume and AAV delivery and transduction was studied under various microbubble size distributions. Similar to our previous studies with a geometrically focused FUS transducer [39], the BBB opening volume increased with bigger microbubbles. The 3-5 μm microbubbles better match the diameter of the capillary vessels, which in mice is on average approximately 6.6 μm [40], thus being able to more effectively deliver a mechanical force on the capillary walls compared to smaller microbubbles. Since the number of microbubbles was controlled across groups, the total injected gas volume was significantly larger for the 3-5 μm and polydisperse groups compared to the 1-2 μm group. Notably, the BBB opening and GFP expression increase as the volume percentage of 3-5 μm bubbles increases (Fig **1J**), which points to the importance of this microbubble size to the effectiveness of ThUS-BBBO. To elucidate whether the smaller BBBO volumes and AAV delivery observed in the 1-2 μm group was due to the lower gas volume injected, future studies will involve normalization of microbubble dose based on volume, rather than number, concentration.

Finally, the AAV serotype was shown to play a crucial role in the transduction efficiency, as well as the types of cells transduced in the brain. The most effective serotype for brain cell transduction was shown to be AAV9, which by far outperformed the AAV5 and AAV2 serotypes (Fig **8**). However, AAV9 also induced significant peripheral organ transduction (Fig **S4D**), and was not specific to a particular type of cells in the brain, whereas AAV5 and AAV2 were more selective with neuronal cells. Additionally, increasing the dose of AAV9 and AAV5 also increased the number of cells expressing GFP in the targeted areas (Fig **8E**), without inducing any observable adverse effects on the mice. Capsid engineering can further improve brain-specific delivery while limiting peripheral organ transduction, and such engineered AAVs will be delivered in future studies.

## Conclusions

In this study, the flow phantom and *in vivo* microbubble activity under ultra-short ThUS pulses was characterized. Short pulse lengths revealed an increased broadband signal in the frequency domain whereas longer pulses exhibited strong and distinct harmonic peaks. Despite the broadband cavitation dominance in short pulses, a smaller extent of microbubble collapse was observed in short pulses compared to long pulses. In the *in vivo* studies, longer pulses also exhibited higher microbubble collapse, which translated to increased instances of microhemorrhage but also increased BBB opening volumes and AAV delivery. PCI-derived cavitation dose was shown to be the most reliable predictor of BBB opening volume as well as AAV transduction for different microbubble sizes as well as different burst sequence parameters. Finally, the transduction efficiency was studied with different AAV serotypes and injection doses, with AAV9 achieving the highest brain and peripheral organ transgene expression. Overall, this study reports on the characterization of microbubble activity associated with ThUS-mediated gene delivery.

## Materials and Methods

### Theranostic Ultrasound system

In this study, the theranostic ultrasound (ThUS) system previously developed by our group [31], [32] was used. A repurposed phased array (P4-1, ATL Philips, 96 Elements, Bandwidth: 1.5-4.0 MHz, Center frequency: 2.5 MHz) was driven at a frequency of 1.5 MHz by a Verasonics research ultrasound system (Vantage 256, Verasonics Inc., Kirkland, WA, USA). A custom MATLAB script was used to both generate ultra-short focused transmits and simultaneously reconstruct the received signal through power cavitation imaging (PCI) as previously described [41]. The ultrasound beam was focused at an axial distance of 35 mm, forming a –6 dB focal region with dimensions 1.5 mm, 8.4 mm, and 14 mm in the lateral, elevational, and axial directions respectively (Fig. **1E-F**), as measured in a water tank using a capsule hydrophone (HGL-0200, Onda Corporation, Sunnyvale, CA, USA). The theranostic pulse sequence used for the phantom study comprised of bursts of 100 ultra-short focused transmits emitted at a pulse repetition frequency (PRF) of 1000 Hz, with bursts repeated at a burst repetition frequency (BRF) of 0.5 Hz. In the phantom study, the cavitation of microbubbles in response to pulse lengths of 1.5, 5, 10 and 20 cycles and a peak-negative pressure of 1.00 MPa was investigated. For the *in vivo* studies, two sequences were used: A single-focus sequence as described above to detect cavitation activity using a PCD with either 1.5 or 10 cycle pulses, and the rapid alternating steering angles (RASTA) sequence (31) to deliver AAV bilaterally to both hemispheres with a pulse length of 1.5 cycles on the left and 10 cycles on the right. The RASTA sequence employed electronic beam steering to alternate between the two hemispheres, applying the odd pulses on the right hemisphere and the even pulses on the left hemisphere.

### Microbubbles

Lipid-shelled microbubbles (MB) were used for all experiments in this study and were synthesized in-house according to a previously published protocol [42]. Freshly opened 1,2-distearoyl-sn-glycero-3-phosphocholine (DSPC, Avanti Polar Lipids Inc., Alabaster, AL, USA) powder and 1,2-distearoyl-sn-glycero-3-phosphoethanolamine-N-[methoxy(polyethylene glycol)-2000] (DSPE-PEG2000, Avanti Polar Lipids Inc., Alabaster, AL, USA) powder were mixed at a 9:1 molar ratio in a glass container with 100 mL of solution containing 80% v/v 1X Phosphate-Buffered Saline (PBS), 10% v/v glycerol, and 10% v/v propylene glycol. The container was submerged in a water bath (Branson Ultrasonic Cleaner 1500, Branson Ultrasonics, Brookfield, CT, USA) that maintained a temperature above 60°C and was continuously sonicated for at 1-3 hours until the lipid powders were fully dissolved, and the solution had a translucent appearance. The lipids were further dissolved using a tip sonicator (Branson Digital Sonifier, Branson Ultrasonics, Brookfield, CT, USA) for 3 minutes at 20% amplitude to ensure complete dilution of the lipids. The lipid mixture was then cooled in an ice bath to room temperature, and two separate procedures were subsequently followed for polydisperse or size-isolated microbubbles. In the phantom studies, polydisperse microbubbles were used, whereas in the *in vivo* studies, both polydisperse and size-isolated microbubbles were injected.

For polydisperse microbubbles, the lipid solution was aliquoted in portions of 1.5 mL into 3 mL glass vials which were stored at 4°C. Immediately before each experiment, a vial was selected and the air in the interstitial space was replaced by perflurobutane (PFB) gas through a series of 5 20-second vacuum and gas cycles. Subsequently microbubbles were formed by vigorously mixing the lipids and the gas in a modified amalgamator (VialMix^TM^, Lantheus Medical Imaging, N. Billerica, MA, USA) for 45 seconds. Following activation, the microbubble size distribution and concentration was quantified using a multisizer (Multisizer 4e Coulter Counter, Beckman Coulter, Indianapolis, IN, USA) by mixing 2 μL of microbubble solution in 10 mL of isoton II (Beckman Coulter™ Isoton™ II Diluent, Beckman Coulter, Indianapolis, IN, USA). On average, the number concentration of microbubbles in the size range 1-10 μm was 1.16 x10^10^ MB/mL, the average size was 1.19 μm, the median size was 0.97 μm and the standard deviation was 0.71 μm (Fig. **1I**).

For size-isolated microbubbles, the tip sonicator was placed on the surface of the lipid solution while PFB gas was being supplied inside the jar. The sonicator was activated for 10 seconds at 100% amplitude to form the microbubbles. After activation, the microbubbles were drawn using a 30 mL syringe and placed in a centrifuge set at 300 x g for 3 minutes, which aimed to separate the formed microbubbles from undissolved lipids and nanobubbles. The separated lipid solution was discarded, and the microbubbles were kept in the syringe, where the volume was replenished to 30 mL using filtered 1X PBS. The remaining MBs were then size-isolated with a series of centrifugation steps. The volume of liquid in the syringe was filled to 30 mL after each centrifugation step. Two groups of size-isolated microbubbles were studied: 1-2 μm, and 3-5 μm. For the first group, the MBs smaller than 2 μm were separated by centrifugation at 220 x g for 1 minute, following by centrifugation of the separated bubble solution at 300 x g for 3 minutes to further discard any bubbles less than 1 μm. For the second group, the bubbles smaller than 3 μm were first removed by centrifugation at 120 x g for 1 minute, and this step was repeated at least 4 times until the separated solution was almost clear. The bubbles remaining in the syringe, containing mainly bubbles larger than 3 μm, were centrifuged at 70 x g for 1 minute once, to separate the bubbles larger than 5 μm, keeping only the 3-5 μm MBs. The size distribution and concentrations of microbubbles was quantified using a multisizer (Fig. **1I-J**).

### Microbubble Flow Phantom Studies

The microbubble cavitation activity under ultra-short ThUS pulses was first studied in a flow phantom (Fig. **1A**). A large glass tank was filled with deionized water and degassed for at least two hours using a portable degassing system (Sonic Concepts, Bothell, WA, USA). The flow channel was made using a 5 cm piece of regenerated cellulose hollow microdialysis fibers (Inner Diameter: 200 μm, Outer Diameter: 216 μm, Spectra/Por, Spectrum Laboratories, Rancho Dominguez, CA) which was attached with glue between two pieces of silicone tubing (Inner Diameter: 500 μm, Outer Diameter: 940 μm, Item 51845K66, McMaster-Carr, Elmhurst, IL). The channel was fixed between two 3D printed poles and placed inside the water tank so that the cellulose tube portion was positioned parallel to the table and the ends of the silicone tubing extended outside the tank. A rubber ultrasound absorber was placed under the flow channel at an approximately 30° angle to divert reflections from the bottom of the tank. Baseline measurements were taken by injecting degassed, deionized water through the phantom, and microbubble cavitation was measured by flowing a solution of polydisperse lipid microbubbles in degassed, deionized water at a concentration of approximately 1 x10^9^ MB/mL. The flow rate was controlled using an infusion pump (Genie Plus, Kent Scientific, Torrington, CT, USA), which pushed the water or microbubble solution through a 1 mL syringe into the flow channel at flow rates varying from 100 to 400 μL/min.

The cavitation activity in the flow channel was monitored using a single element transducer (C384-SU, Olympus, Waltham, MA, Center frequency: 3.5 MHz, –6dB Bandwidth: 2.30-4.62 MHz) as a passive cavitation detector (PCD). The PCD was placed in the water tank and positioned parallel to the table and directly across the cellulose tube (Fig. **1A**). The transducer was connected to a pulser/receiver (5072PR, Olympus, Waltham, MA) which gave the signal a 20dB gain, and which was in turn connected to a computer oscilloscope (Picoscope 5000 Series, Pico Technology, St. Neots, UK). Cavitation data were collected using the Picoscope software at a sampling rate of 62.5 MHz for 50 ms per acquisition, and the data were saved as MATLAB-4 files (.mat) for further processing.

The P4-1 transducer was fixed on a 3-axis positioning system (VXM, Velmex Inc., Bloomfield, NY, USA) and B-mode imaging at 2.5 MHz was used to focus the array at an axial distance of 35 mm over the cellulose tube (Fig. **1G-H**). The lateral alignment of the transducer with the PCD was achieved by transmitting focused pulses with the P4-1 and translating the array laterally until no further increase in the received signal on the PCD was observed. The same method was finally used for the elevational alignment of the P4-1 with the cellulose tube. The MB cavitation activity was recorded for 5 bursts per pulse length and flow rate combination. A summary of the experimental parameters is given in Table **1**.

**Table 1.**
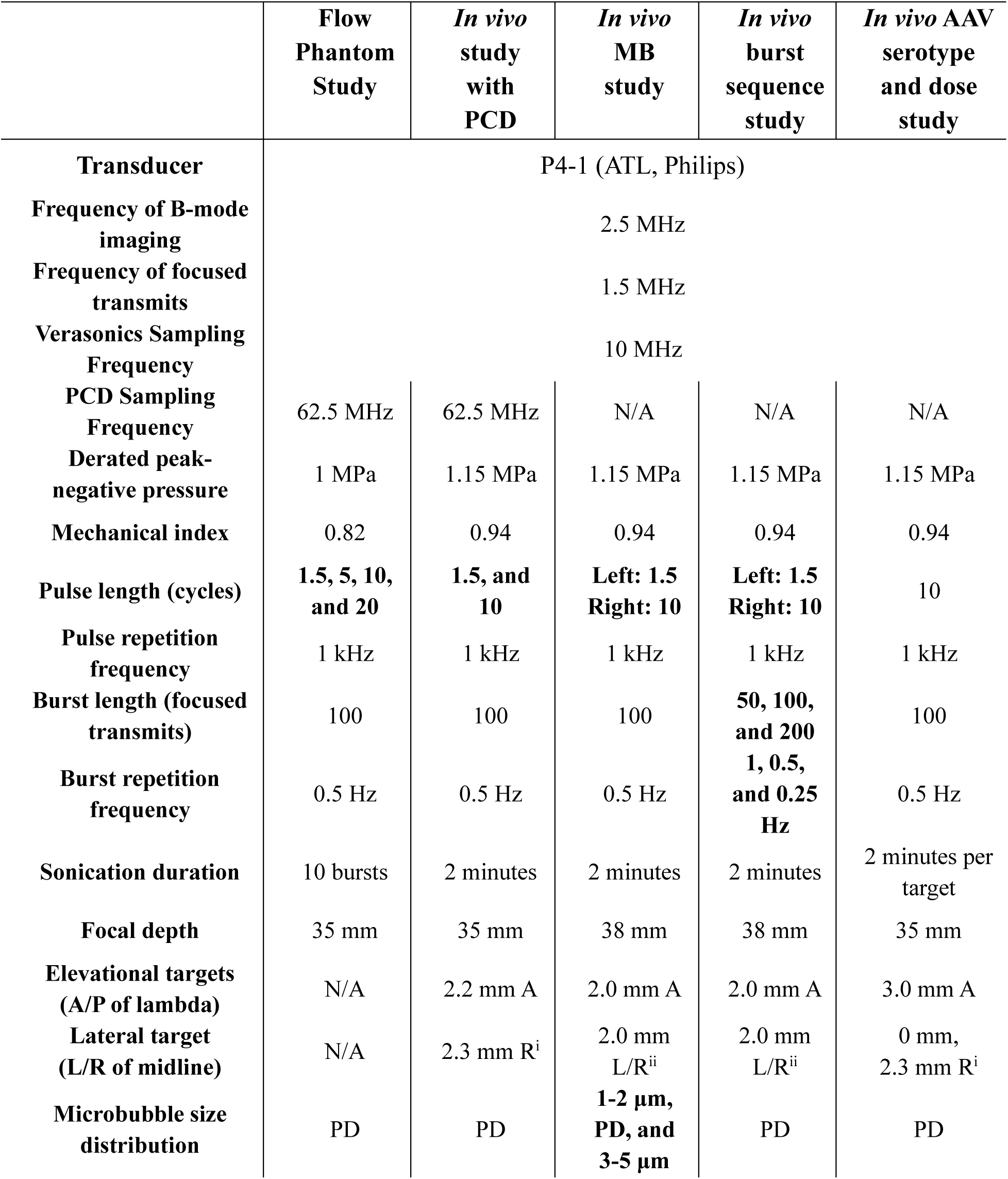
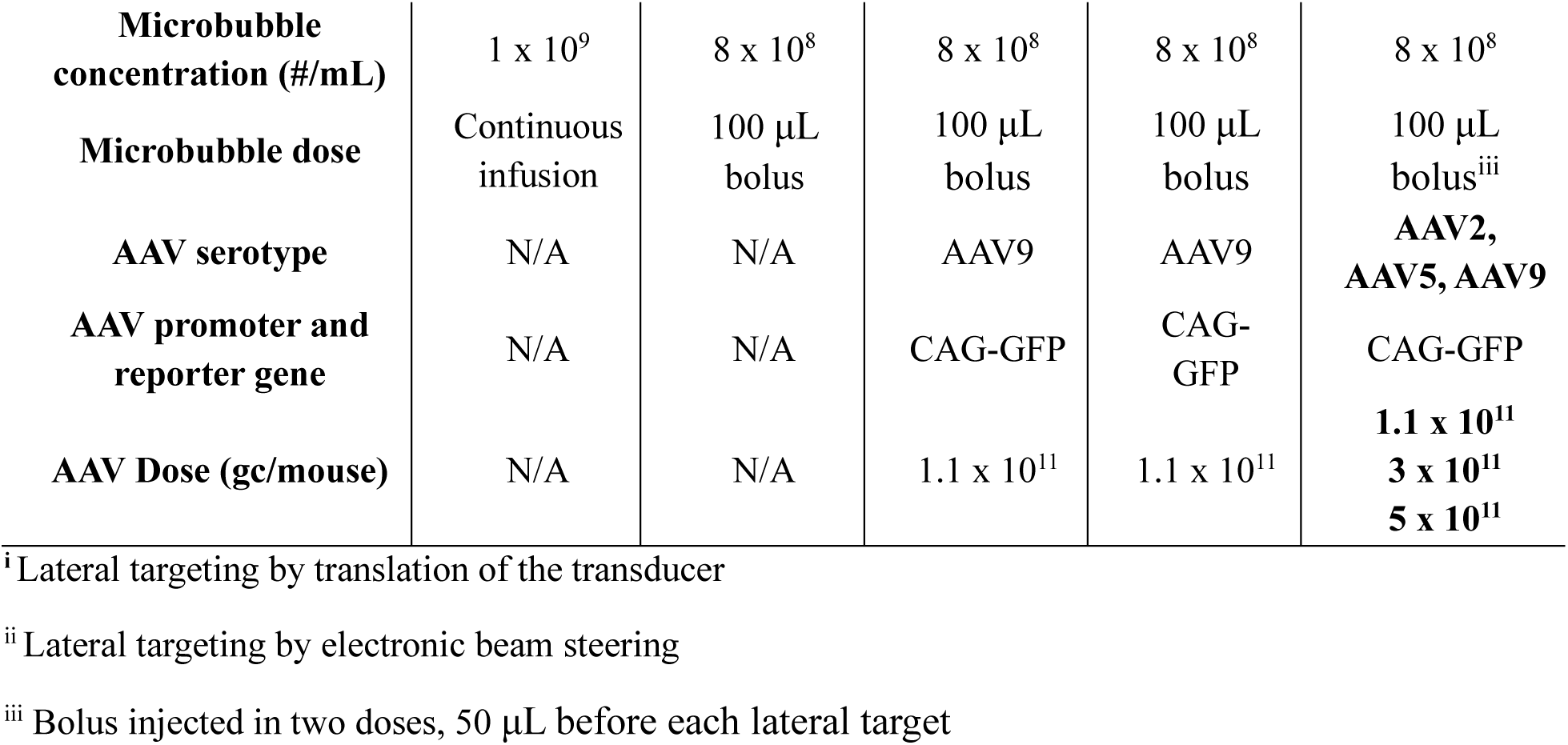
Experimental parameters for the flow phantom and *in vivo* experiments. Values in boldface represent a study variable. A/P: Anterior-posterior; GC: Gene copies; L/R: Left-right PCD: Passive Cavitation Detector; PD: Polydisperse

**Table 2.**
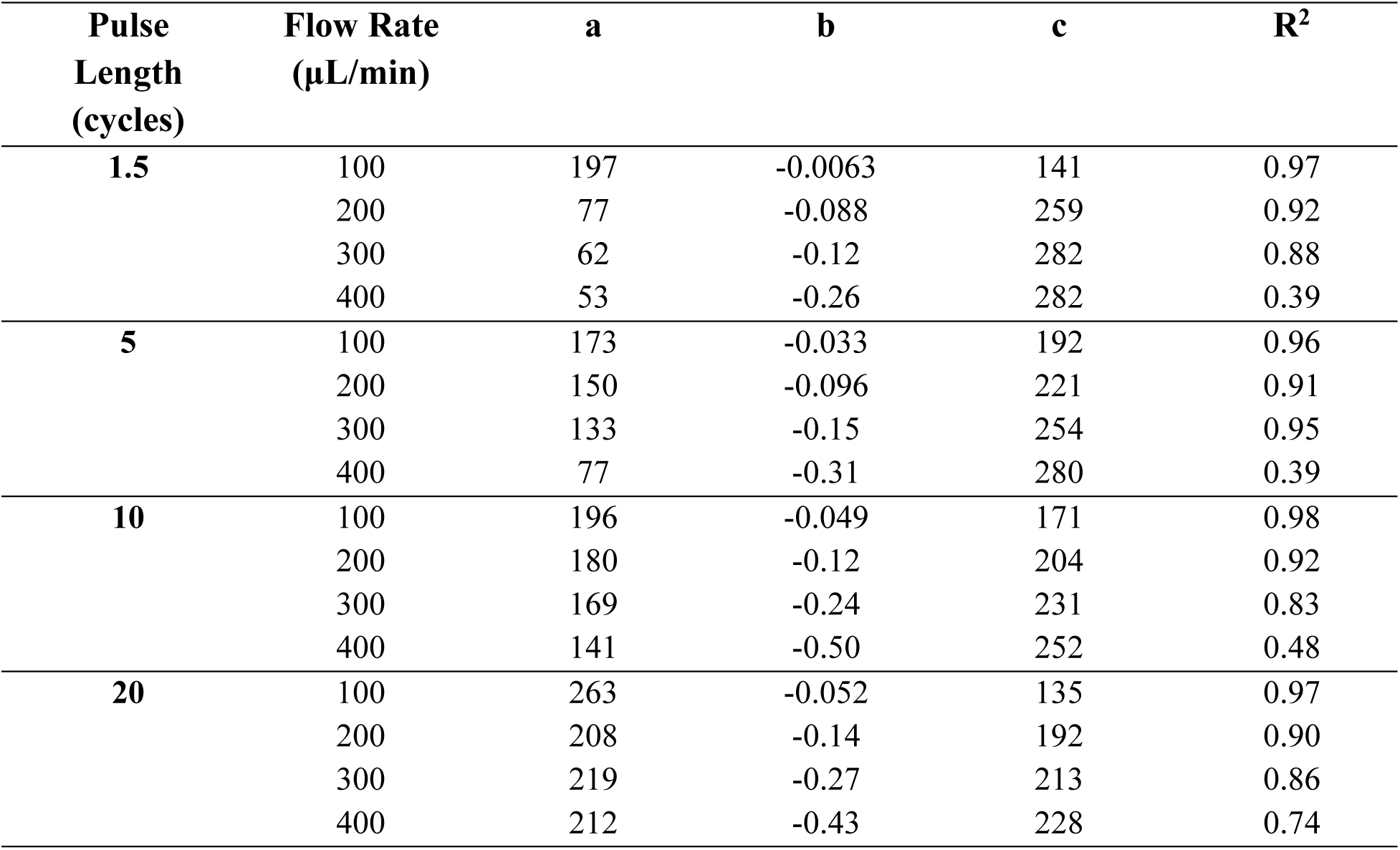
Exponential fit parameters for variable pulse lengths and flow rates in the flow phantom experiment. An exponential decay model of the form ***y*** = ***a*** ∗ ***exp***(***b*** ∗ ***x***) + ***x*** was fitted to the time domain data. The goodness of fit was evaluated by the R^2^ metric.

| Pulse Length (cycles) | Flow Rate ($\mu\text{L}/\text{min}$) | a | b | c | $R^2$ |
| --- | --- | --- | --- | --- | --- |
| 1.5 | 100 | 197 | -0.0063 | 141 | 0.97 |
|  | 200 | 77 | -0.088 | 259 | 0.92 |
|  | 300 | 62 | -0.12 | 282 | 0.88 |
|  | 400 | 53 | -0.26 | 282 | 0.39 |
| 5 | 100 | 173 | -0.033 | 192 | 0.96 |
|  | 200 | 150 | -0.096 | 221 | 0.91 |
|  | 300 | 133 | -0.15 | 254 | 0.95 |
|  | 400 | 77 | -0.31 | 280 | 0.39 |
| 10 | 100 | 196 | -0.049 | 171 | 0.98 |
|  | 200 | 180 | -0.12 | 204 | 0.92 |
|  | 300 | 169 | -0.24 | 231 | 0.83 |
|  | 400 | 141 | -0.50 | 252 | 0.48 |
| 20 | 100 | 263 | -0.052 | 135 | 0.97 |
|  | 200 | 208 | -0.14 | 192 | 0.90 |
|  | 300 | 219 | -0.27 | 213 | 0.86 |
|  | 400 | 212 | -0.43 | 228 | 0.74 |

### Cavitation Dose Calculation

The time-domain data were imported to MATLAB and the burst signal was separated from pre– and post-acquisition noise by applying a rectangular window of 100 ms duration. The burst was further divided into individual pulses and another rectangular window was applied to isolate the pulse signal from reflections and noise. The isolated pulses were transformed to the frequency domain using MATLAB’s FFT function with 2^19^ points, which were deemed sufficient as they far exceeded the number of time-domain points and allowed for adequate zero-padding. The total cavitation dose (CD) was first calculated for frequencies between 3.5 MHz and 14 MHz:

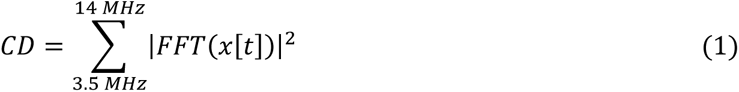

where *x*[*t*] is the time-domain amplitude of the received signal from the PCD, and FFT is the fast Fourier transform. Harmonic and ultraharmonic cavitation doses were subsequently quantified by summation of the FFT values within 200 kHz bands centered around the harmonic and ultraharmonic frequencies. For the PCD data, the third to ninth harmonic (*harm*) and ultraharmonic (*ultra*) frequencies were selected. The fundamental and second harmonic and ultraharmonic frequencies were not included in the calculation of cavitation dose to avoid contamination of the secondary microbubble signal from reflections originating from the ThUS beam and the skull.

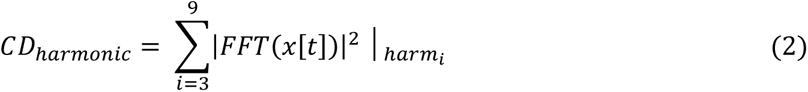

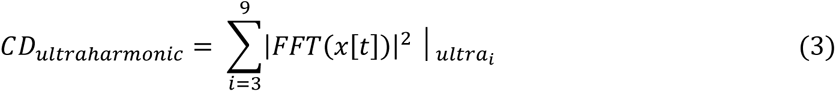

Finally, the contributions outside the harmonic and ultraharmonic frequency bands were considered to be “broadband cavitation”.

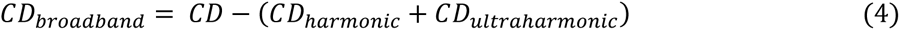

All cavitation components were calculated for each individual pulse in a single burst, and all the pulse CDs were summed to yield the CD of a burst. For baseline measurements, the CD of all bursts was averaged to obtain the mean baseline cavitation for each condition, which was then used to calculate the normalized microbubble cavitation dose.

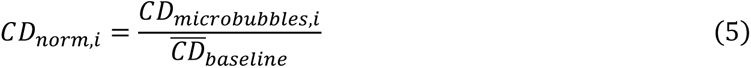

For the calculation of the cavitation dose from the data received by the P4-1, the raw radio frequency (RF) data from each element were transformed to the frequency domain using MATLAB’s FFT function, before being summed across all elements to obtain the amplitude-only frequency spectrum. Since the sampling frequency of the Vantage system was 10 MHz, analysis of frequencies up to 5 MHz was feasible, thus restricting the calculation of the cavitation dose. For this reason, we modified the total cavitation dose to include frequencies from 2.5 MHz to 5 MHz, thus considering both the second and third harmonic and ultraharmonic frequencies but excluding the fundamental frequency. Harmonic and ultraharmonic cavitation doses were calculated by summation of all frequency bins within 200 kHz bands centered around the respective frequencies.

### Power Cavitation Imaging

In addition to the frequency domain cavitation calculation described above, the cavitation dose in *in vivo* experiments was calculated using the power cavitation imaging (PCI) maps that were reconstructed after every ThUS burst as previously described [31], [41]. Briefly, the RF data after each ultra-short ThUS pulse were passively received using the same P4-1 phased array that was used for the focused transmits. The RF data were beamformed using a GPU-accelerated delay-and-sum (DAS) algorithm, and a spatiotemporal clutter filter using singular value decomposition (SVD) was applied to reduce skull signal contributions by discarding the eigenvectors corresponding to singular values below 8. The individual-pulse PCI maps from an entire burst were summed to create a high-contrast cavitation map. The PCI-derived cavitation dose was calculated by summing the pixels of each PCI frame throughout the sonication.

### *In Vivo* ThUS-BBBO and AAV Delivery

All animal experiments were reviewed and approved by the Institutional Animal Care and Use Committee at Columbia University. Four *in vivo* studies were carried out to characterize ThUS cavitation and its effect on BBB opening and AAV delivery. In the first study, the transcranial cavitation dose of ThUS in a mouse model was evaluated using a PCD. Two 8-week-old male C57BL/6J mice were subjected to ThUS-mediated BBBO (ThUS-BBBO) as previously described, unilaterally on the right hippocampus and using two ThUS pulse lengths: 1.5, and 10 cycles. The ThUS pulse sequence comprised of 100 focused pulses, repeated at a PRF of 1 kHz, and a BRF of 0.5 Hz, and a derated peak rarefactional pressure of 1.15 MPa was applied. A higher pressure than the flow phantom experiments was chosen to assess MB cavitation in conditions similar to our previous gene editing study [43]. The cavitation was monitored using a separate single-element transducer (V312-SU, Olympus, Waltham, MA, Center frequency: 10 MHz, –6dB Bandwidth: 5.74-12.42 MHz) acting as a PCD, which was connected to a pulser receiver (5072PR, Olympus, Waltham, MA) and a computer oscilloscope (Picoscope 5000 Series, Pico Technology, St. Neots, UK). The PCD was placed at an angle in front of the P4-1 so that the field of view of the PCD aligned with the focus of the theranostic array. Alignment of the PCD to the focus of the P4-1 was confirmed using a flow phantom with microbubbles immediately prior to the animal experiments. A schematic of this setup is shown in Figure **1B**.

In the second study, the cavitation, BBBO volume and AAV delivery were evaluated under variable microbubble size distributions. Eight-week-old male C57BL/6J mice were randomly divided in groups of 6, with each group receiving one of three microbubble formulations: 1-2 μm, 3-5 μm size-isolated microbubbles, or polydisperse microbubbles. The size distribution of the three microbubble groups is shown in Figures **1I-J**. The RASTA sequence [32] was applied to bilaterally induce ThUS-BBBO, with alternating 1.5-cycle pulses on the left and 10-cycle pulses on the right hemisphere, and a total of 100 focused transmits repeated at a PRF of 1 kHz and a BRF of 0.5 Hz. The derated peak-negative pressure was 1.15 MPa.

In the third study, the effect of the number of pulses per burst, or burst length (BL), as well as the burst repetition frequency (BRF), on AAV delivery and BBBO volume was assessed. Similarly to the second study, three groups of 8-week-old male C57BL/6J mice were randomly formed and each group received one of three combinations of BL and BRF: 200 pulses with 0.25 Hz BRF, 100 pulses with 0.5 Hz BRF, or 50 pulses with 1 Hz BRF. Polydisperse microbubbles were used for all groups; therefore, the group receiving 100 pulses with 0.5 Hz BRF was receiving the same parameters as the group receiving polydisperse microbubbles in the second study. The same group of mice was therefore used for both studies, to adhere to the 3Rs principle. All other ThUS parameters were kept the same as the second study. A schematic of the experimental setup for the second and third studies is shown in Figure **1C**.

The last study focused on the transgene expression efficiency in the brain using the same ThUS parameters but different AAV serotypes and injection doses. A single focus sequence with BL of 100 pulses and a BRF of 0.5 Hz was used, while polydisperse microbubbles were injected with the AAV constructs. The AAV9-CAG-GFP construct (Addgene viral prep 37825-AAV9) used in the previous two studies was administered in doses of 3 x 10^11^, and 5 x 10^11^ gc/mouse, the AAV5-CAG-GFP construct (Addgene viral prep 37825-AAV5) was administered in doses of 1.1 x 10^11^, and 3 x 10^11^ gc/mouse, and the AAV2-CAG-GFP construct (Addgene viral prep 37825-AAV2) was injected at a dose of 3 x 10^11^ gc/mouse. In this study, only one mouse per condition was used.

The mice were anesthetized with a mixture of oxygen and 1-2% isoflurane until no reflex to the toe pinch test was visible. Their heads were shaved, depilated, and placed in a stereotaxic apparatus. Degassed ultrasound gel was applied on their depilated heads and a water bath with degassed and deionized water was placed on top of their heads. The lambdoid suture was identified through the scalp and a metallic crosshair was positioned on top of it. The P4-1 transducer was lowered inside the water bath and B-mode imaging at 2.5 MHz was used to guide the transducer over the head, and using anatomical features and the reflection from the metallic crosshair, the focus was targeted over the lambdoid suture. In the PCD study, to target the right hippocampus, the transducer was translated by 2.2 mm anteriorly and 2.3 mm to the right, and the beam was axially focused at 35 mm. For the microbubble and burst sequence studies, the transducer was translated elevationally by 2 mm, and electronic lateral beam steering 2 mm right and left was applied. Additionally, a deeper axial focus was applied, at 38 mm, to target the substantia nigra. Finally, in the AAV serotype and dose study, the transducer was translated by 3 mm anteriorly, and two lateral targets, one over the center of the brain and one 2.3 mm to the right were sonicated. A tail-vein catheter was placed in the mice’s depilated tails, and a small amount of sterile saline was injected while ThUS pulses were applied, to obtain baseline cavitation signal. Following baseline acquisition, the microbubble, or microbubble + AAV solutions were injected as a single 100 μL bolus through the tail vein. Two minutes after injection the mice were removed from the stereotaxic apparatus and placed in cages until MRI was carried out for BBB opening confirmation. A summary of the experimental parameters is given in Table **1**.

### Magnetic Resonance Imaging

BBB opening and safety were confirmed within two hours after sonication using contrast-enhanced T1-weighted MRI and T2*-weighted MRI, respectively. Before administration of the contrast agent, the mice were anesthetized with a mixture of oxygen and 1-2% isoflurane and then placed inside a 9.4 T vertical bore system (Ascend 400WB, Bruker, Billerica, MA, USA) where a T2*-weighted 2D FLASH sequence (TR: 350 ms, TE: 12 ms, flip angle: 40°, number of excitations: 20, field of view 20 mm × 18 mm) was carried out to assess the presence of hemorrhage. The mice were then intraperitoneally injected with 0.2 mL of gadodiamide solution (Omniscan^TM^, GE Healthcare, Princeton, NJ) and a T1-weighted 2D FLASH sequence (TR: 230 ms, TE: 3.3 ms, flip angle: 70°, number of excitations: 6, field of view: 25.6 mm × 25.6 mm) was performed to confirm BBB opening.

### Histological Preparation and Microscopy

Three weeks after the ThUS-BBBO procedure the mice were euthanized for immunohistochemical analysis. The mice were heavily anesthetized with a mixture of oxygen and 3% isoflurane, and after anesthesia was confirmed, they were exsanguinated and transcardially perfused with cold, USP-grade 1X PBS for at least 5 minutes until the liver changed color from deep red to light brown, followed by 4% paraformaldehyde for at least 5 minutes. The brains were then extracted from the skull and placed in a solution of 4% paraformaldehyde for 24 hours, followed by 30% sucrose in 1X PBS for at least 24 hours in preparation for sectioning.

Coronal sections were obtained using a cryostat (Leica CM1850, Leica Biosystems, Buffalo Grove, IL, USA) or a vibratome (Leica VT100S, Leica Biosystems, Buffalo Grove, IL, USA) at a thickness of 40 μm. Sections corresponding to the regions of BBB opening were selected for immunofluorescence staining. After being washed in 1X Tris-buffered saline (TBS) three times for 10 minutes, the sections were incubated for 60 minutes in a blocking solution containing 1X TBS, 0.3% TritonX-100 and 4% normal donkey serum (Jackson ImmunoResearch Inc., West Grove, PA, USA). The sections were then transferred in the primary antibody solution containing chicken anti-GFP antibody (1:1000 dilution in 1X TBS, 0.3% TritonX-100 and 4% normal horse serum, Novus Biologicals, Centennial, CO, USA), where they were incubated at 4°C in the dark for 48 hours. After incubation, the sections were washed twice for 10 minutes in 1X TBS, and once for 30 minutes in 1X TBS, 0.3% TritonX-100 and 4% normal horse serum, and were subsequently incubated for 2 hours in the dark in the secondary antibody solution containing donkey anti-chicken Alexa Fluor 488 (1:250 dilution in 1X TBS, 0.3% TritonX-100 and 4% normal horse serum, Invitrogen, Waltham, MA, USA). Finally, the sections were washed thrice for 10 minutes in 1X TBS, before being mounted on glass microscopy slides using DAPI-containing mounting medium (ab104139, Abcam, Cambridge, United Kingdom).

For safety assessment, one mouse from each group of the microbubble and burst sequence studies was euthanized at 24 hours post-ThUS-BBBO. After tissue fixation as described above, the brains were transferred to 1X PBS and were paraffin-embedded for coronal sectioning in 5 μm thick sections. Hematoxylin and Eosin (H&E) staining was then carried out to assess cellular damage and microhemorrhage.

The GFP-stained sections were viewed under a fluorescent microscope (Olympus BX61, Evident Corporation, Tokyo, Japan), and the H&E sections were in a bright field digital slide scanner (Leica Aperio AT2 DX, Leica Microsystems Inc., Buffalo Grove, IL, USA) to visualize the expression of GFP and the presence of microhemorrhage respectively. GFP expression on stained sections was quantified using a customized MATLAB script.

### Statistical Analysis

Statistical analysis was conducted on GraphPad Prism (GraphPad Software Inc., La Jolla, CA). For the *in vivo* experiments, linear regression analysis was carried out to correlate the cavitation dose with BBBO volume and R^2^– and p-values were obtained to assess the goodness of fit and significance of the analysis. Additionally, the differences in GFP expression between groups was assessed using one-way ANOVA, with post-hoc Tukey’s multiple comparison tests. Statistical significance was set for p<0.05.

## Abbreviations

AAV: Adeno-associated virus
BBB: Blood-brain barrier
BBBO: Blood-brain barrier opening
BRF: Burst repetition frequency
CD: Cavitation dose
CNS: Central nervous system
DFT: Discrete Fourier Transform
FFT: Fast Fourier Transform
FT: Focused transmit
FUS: Focused ultrasound
FUS-BBBO: Focused ultrasound-mediated blood-brain barrier opening
GC: Gene copies
GFP: Green fluorescent protein
H&E: Hematoxylin and eosin
MB: Microbubble
MI: Mechanical index
MRI: Magnetic resonance imaging
RASTA: Rapid alternating steering angles
RF: Radio frequency
PBS: Phosphate-buffered saline
PCD: Passive cavitation detector
PCI: Power cavitation imaging
PFB: Perflurobutane
PRF: Pulse repetition frequency
SVD: Single value decomposition
TBS: Tris-buffered saline
ThUS: Theranostic ultrasound
ThUS-BBBO: Theranostic ultrasound-mediated blood-brain barrier opening
USP: United States pharmacopeia.

## Acknowledgements

This study was funded in part by the National Institutes of Health under grants R01AG038961, R01EB009041, and R56AG038961, the Defense Advanced Research Projects Agency (DARPA) under Contract No. N66001-19-C-4020, and the Focused Ultrasound Foundation. FNT was also supported by the Onassis Foundation under contract number F ZT 072-1/2023-2024. The authors wish to thank UEIL members Rebecca L. Noel, Ph.D., Nancy Kwon, M.S., Sergio Jiménez-Gambín, Ph.D., Aparna Singh, Ph.D., Yangpei Liu, M.S., Parth Gami, M.S., Moshe J. Wilner M.S., and Melody DiBenedetto B.S., as well as interns Hadrien D. Padilla, Rashell K. Ramirez, Maya Yie, Chloe Lugg, Red Nochomovitz, Claire Comisarow and Chloe Comisarow for their support and fruitful scientific discussions. The authors also wish to acknowledge the Columbia University Medical Center Molecular Pathology/MSPR core for histological services. The views, opinions, and/or findings expressed in this study are those of the authors and should not be interpreted as representing the official views or policies of the Department of Defense or the U.S. Government. Some figures were created using Biorender.com.

## Contributions

FNT, AJB, and EEK designed the study and the methodology. FNT synthesized and isolated microbubbles. FNT, CL, RJ and SB conducted the flow phantom experiments. FNT, AJB and DAJ conducted the *in vivo* experiments. FNT, DAJ, AT, and SLG sectioned, stained and imaged immunofluorescent brain sections. FNT processed and analyzed ultrasound, MRI and microscopy data, created figures and tables, drafted and revised the manuscript. AJB, CL, RJ, and SB also contributed to data discussions and review. EEK acquired funding, provided resources for the study and contributed to discussions and reviews of the manuscript.

## Supplementary Material

Please see attached supplementary materials (Fig. S1-S4).

