## Supplementary Materials for "Characterization of Microbubble Cavitation in Theranostic Ultrasound-mediated Blood-Brain Barrier Opening and Gene Delivery"

**Included in this file:**

Figures S1-S4

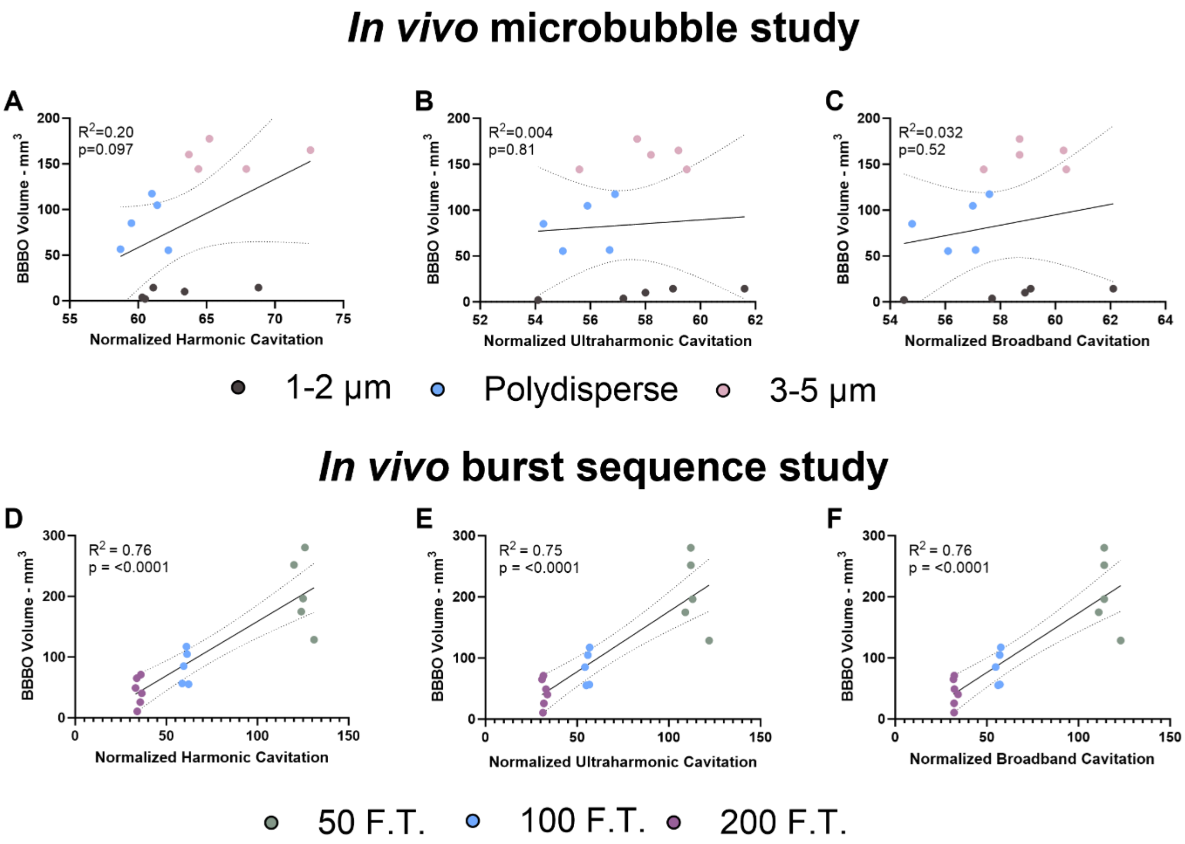


**Figure S1.** Correlation of BBB opening volume with cavitation types calculated in the frequency domain using P4-1 data. **A-C)** Correlation of BBB opening volume with harmonic (A), ultraharmonic (B), and broadband (C) cavitation doses for the *in vivo* microbubble study. There is no good correlation within or between groups for any cavitation type. **D-F)** Correlation of BBB opening volume with harmonic (D), ultraharmonic (E), and broadband (F) cavitation doses for the *in vivo* burst sequence study. Although there is a significant positive correlation across groups for all cavitation types, there is no significant correlation within groups for any cavitation type.

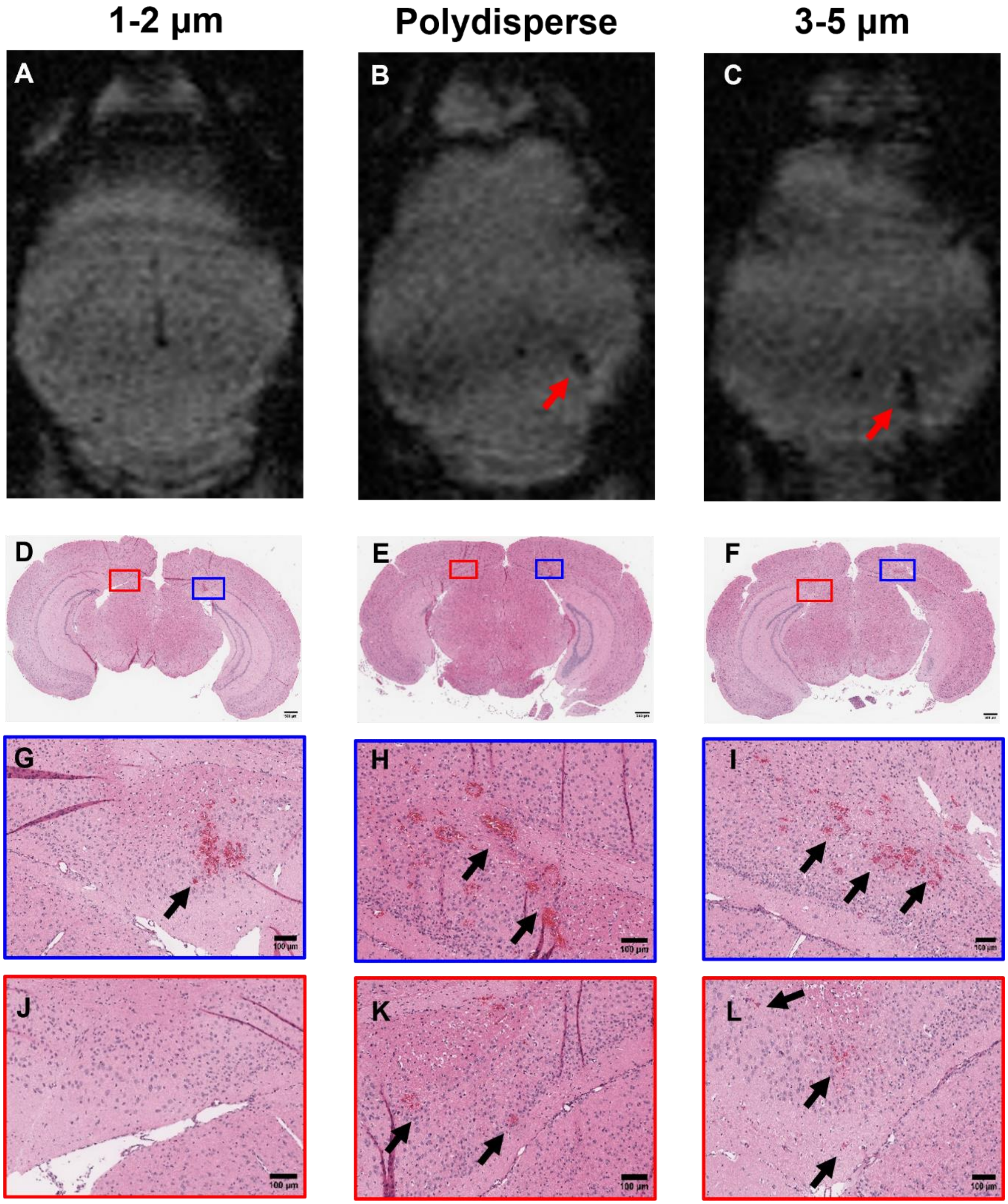


**Figure S2.** Safety assessment of ThUS-BBBO for the *in vivo* microbubble study. **A-C)** Axial T2*-weighted MRI images of mice injected with 1-2 μm (A), polydisperse (B), and 3-5 μm (C) MBs taken on the day of sonication. The presence of hypointense regions, indicated by red arrows in (B) and (C) indicates the presence of hemorrhage on the right hemisphere of mice injected with polydisperse and 3-5 μm MBs, which was sonicated with 10-cycle pulses. **D-F)** Hematoxylin and eosin-stained brain sections of mice injected with 1-2 μm (A), polydisperse (B), and 3-5 μm (C) MBs and sacrificed 1 day after sonication. The scale bars are 500 μm. **G-H)** Magnified regions from the right hemisphere treated with 10 cycles, indicated respectively in (D)-(F) by blue rectangles. Black arrows point to regions where erythrocyte extravasation was observed. Microhemorrhage is present for all conditions. The scale bars are 100 μm. **J-L)** Magnified regions from the left hemisphere treated with 1.5 cycles, indicated respectively in (D)-(F) by red rectangles. Black arrows point to regions where erythrocyte extravasation was observed. Microhemorrhage is present for the polydisperse (K) and 3-5 μm (L) groups but not for the 1-2 μm (J) group, and the erythrocyte extravasation is smaller compared to the right hemisphere. The scale bars are 100 μm.

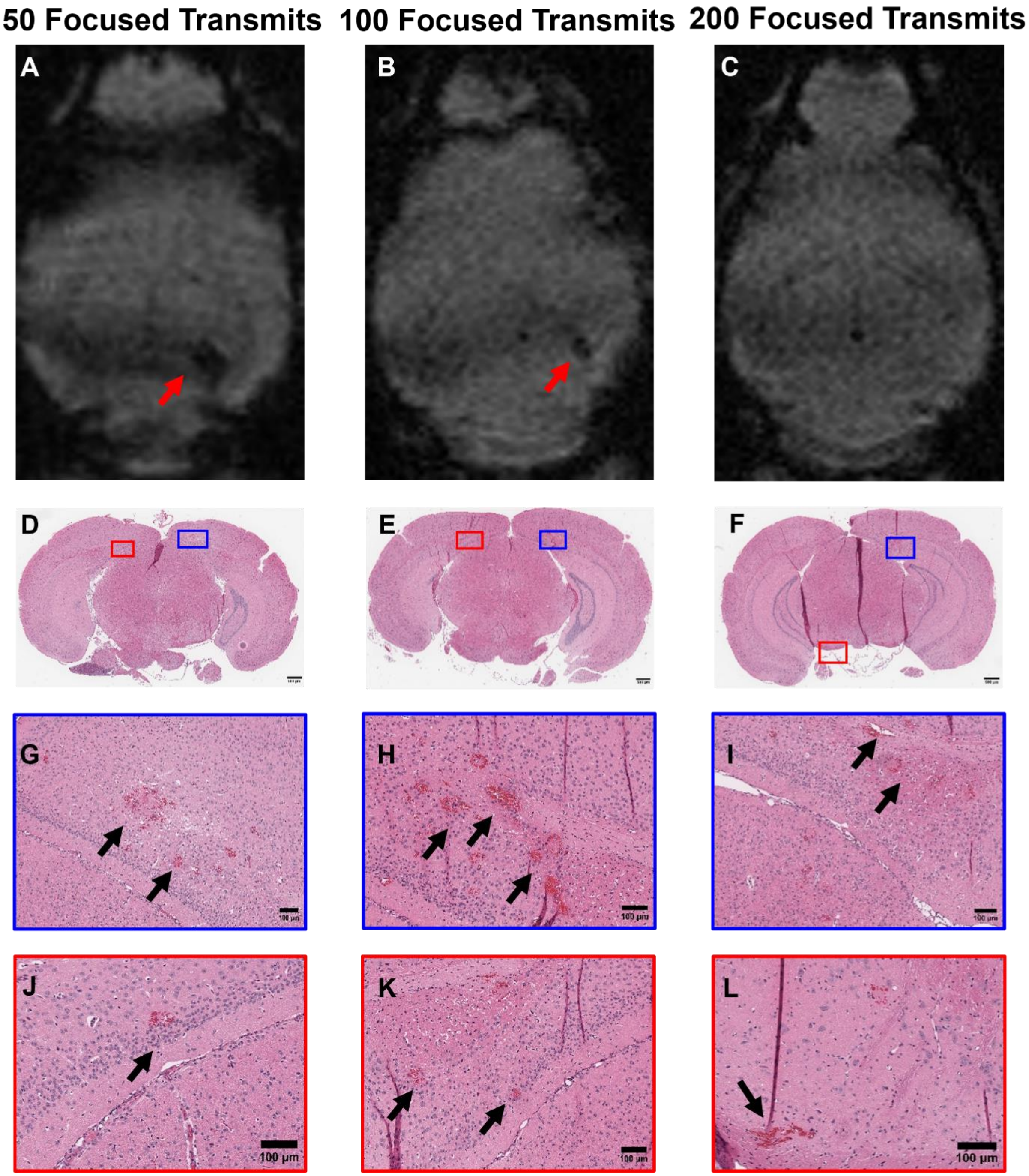


**Figure S3.** Safety assessment of ThUS-BBBO for the *in vivo* burst sequence study. **A-C)** Axial T2*-weighted MRI images of mice sonicated with 50 (A), 100 (B), and 200 (C) focused transmits per burst on the day of sonication. The presence of hypointense regions, indicated by red arrows in (A) and (B) indicates the presence of hemorrhage on the right hemisphere (sonicated with 10-cycle pulses) of mice sonicated with 50 (A) and 100 (B) pulses per burst. **D-F)** Hematoxylin and eosin-stained brain sections of mice sonicated with 50 (D), 100 (E), and 200 (F) focused transmits per burst and sacrificed 1 day after sonication. The scale bars are 500 µm. **G-H)** Magnified regions from the right hemisphere treated with 10 cycles, indicated respectively in (D)-(F) by blue rectangles. Black arrows point to regions where erythrocyte extravasation was observed. Microhemorrhage is present for all conditions. The scale bars are 100 μm. **J-L)** Magnified regions from the left hemisphere treated with 1.5 cycles, indicated respectively in (D)-(F) by red rectangles. Black arrows point to regions where erythrocyte extravasation was observed. Microhemorrhage is present for all conditions, and the erythrocyte extravasation is smaller compared to the right hemisphere. The scale bars are 100 μm.

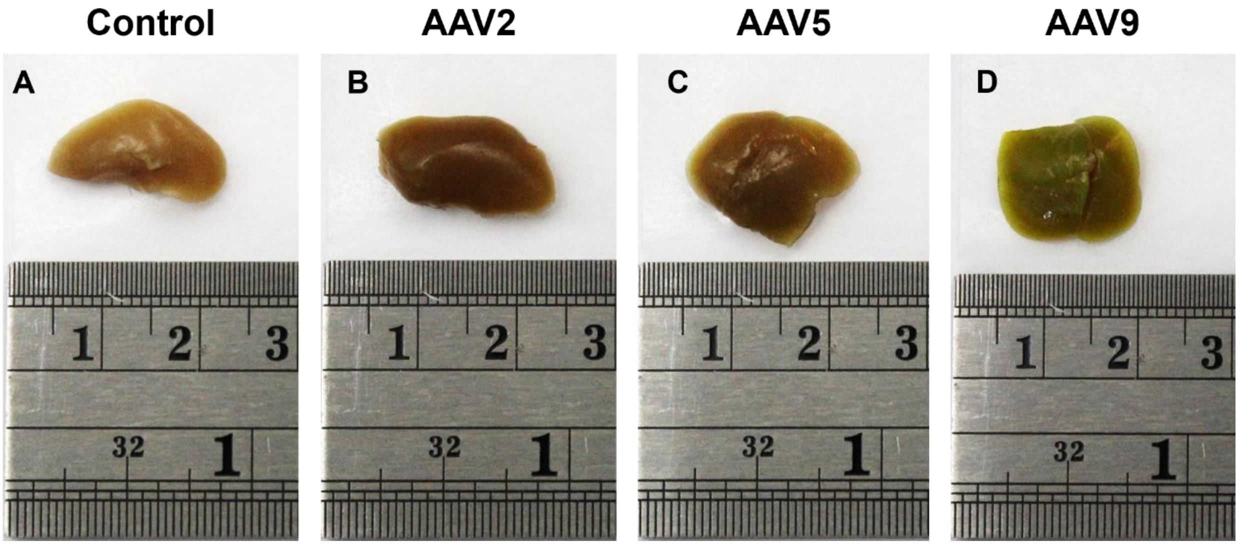


**Figure S4. A-D)** Images of livers post-perfusion and fixation from mice injected with no AAV (A), or a dose of 3 x $10^{11}$ gc/mouse of AAV2 (B), AAV5 (C), and AAV9 (D), all encoding the green fluorescent protein (GFP) using the CAG promoter. The mice injected with AAV were sacrificed 3 weeks after AAV injection. Significant color change is observed in the liver of the mouse injected with AAV9, indicating extensive off-target transgene expression. Ruler: top scale in centimeters, bottom scale in inches.
